# Edited Filamin A in myeloid cells reduces intestinal inflammation and protects from colitis

**DOI:** 10.1101/2025.03.25.643676

**Authors:** Riem Gawish, Rajagopal Varada, Florian Deckert, Anastasiya Hladik, Linda Steinbichl, Laura Cimatti, Katarina Milanovic, Mamta Jain, Natalya Torgasheva, Andrea Tanzer, Kim De Paepe, Tom Van de Wiele, Bela Hausmann, Michaela Lang, Martin Pechhacker, Nahla Ibrahim, Ingrid DeVries, Christine Brostjan, Michael Sixt, Christoph Gasche, Louis Boon, David Berry, Michael F. Jantsch, Fatima C. Pereira, Cornelia Vesely

**Author notes:** Correspondence should be addressed to: Cornelia Vesely, Medical University of Vienna, Center of Anatomy and Cell Biology, Division of Cell and Developmental Biology, Schwarzspanierstrasse 17, A-1090 Vienna, AUSTRIA,; Fatima C. Pereira, University of Vienna, Center for Microbiology and Environmental Systems Science, Department of Microbiology and Ecosystem Science, Djerassiplatz 1, A-1030 Vienna, AUSTRIA,; Riem Gawish, Medical University of Vienna, Department of Medicine I, Research Division of Infection Biology Währinger Gürtel 18-20, A-1090 Vienna, AUSTRIA.

## Abstract

Patho-mechanistic origins and disease dynamics of ulcerative colitis are still poorly understood. The actin-crosslinker Filamin A (FLNA) impacts cellular responses through interaction with cytosolic proteins. FLNA exists in two forms that differ only in one amino acid: genome-encoded FLNA^Q^ and FLNA^R^ - generated by post-transcriptional A-to-I editing. FLNA is edited in fibroblasts, smooth muscle- and endothelial cells in the colon. We identified the FLNA editing status as a key determinant of colitis severity. FLNA editing was highest in healthy colons and reduced during acute murine and human colitis. Mice that exclusively express edited FLNA^R^ and do not downregulate editing upon challenge were highly resistant to DSS-induced colitis, whereas fully unedited FLNA^Q^ animals developed severe inflammation. While the genetic induction of FLNA editing influenced transcriptional states of structural cells and the microbiome composition, we found that FLNA^R^ exerts protection specifically via its influence on myeloid cells, which are not edited under physiological conditions. Introducing fixed, fully edited FLNA^R^ did not hamper normal cell migration but reduced macrophage inflammation and rendered neutrophils less prone to NETosis. In conclusion, loss of FLNA editing correlates with colitis severity, and targeted FLNA editing of myeloid cells might serve as a novel therapeutic approach in intestinal inflammation.

**Summary:** In this study, Gawish et al. show that RNA editing of the actin cross-linker FLNA is similarly regulated in mice and humans and that the targeted induction of edited FLNA^R^ in myeloid cells governs resistance to DSS-induced colitis, revealing its potential in IBD therapy.

## Introduction

The actin cytoskeleton is a complex and dynamic network composed of actin filaments and binding proteins, which define cell morphology and function. Filamin A (FLNA) is a large actin-crosslinking protein, ubiquitously expressed and composed of 24 Ig-like domains. Functionally, FLNA links actin filaments with the cellular cortex and, via Ig-domains 21-23, acts as a scaffolding platform for a variety of proteins, thereby tuning cellular signaling (1). FLNA deficiency in mice is embryonically lethal, causing cardiac defects and abnormal epithelial and endothelial organization (2). In humans, *FLNA* mutations cause a broad range of congenital disorders highlighting its cell type specific functions (3). Interestingly, certain FLNA mutations cause gastrointestinal dysfunction like severe constipation, intestinal pseudo-obstruction and short bowel syndrome. Likewise, FLNA critically influences intestinal development in mice (4). FLNA mRNA is post-transcriptionally modified by adenosine (A) to inosine (I) editing, an RNA modification catalyzed by the adenosine deaminase acting on double stranded RNA 2 (ADAR2). As I is interpreted as guanosine (G) by most cellular machineries, exonic A to I editing can cause protein recoding and proteome diversification. Aside from the physiological functions of A-to-I editing, genetically engineered ADAR enzymes can be redirected to specific targets. After the first demonstration of site-directed RNA editing (SDRE) the potential for therapeutic RNA editing became more and more evident, with recent approaches mainly focusing on the transient and cell modulatory features of SDRE (5, 6).

FLNA recoding is highly conserved in vertebrates and changes a glutamate (Q) to an arginine (R), right in the center of the protein-interaction-scaffolding region of FLNA (Ig-domain 22) (7). Using mice that exclusively express unedited FLNA^Q^ or edited FLNA^R^ (8, 9), we recently showed that edited FLNA^R^ increases cellular stiffness and adhesion in murine fibroblasts, while unedited FLNA^Q^ renders cells more flexible and facilitates cell migration (10). In mice, FLNA is edited in vasculature and accordingly, regulates blood pressure (9) and promotes tumor angiogenesis (8). In the murine gut, FLNA editing is high with 80 - 90 % in the stomach and 60 - 80 % in the large intestine (7, 11). In human colon biopsies editing frequencies up to 26 % have been reported (12). However, the role of FLNA editing in the gut is still unexplored as of today.

The gastrointestinal tract requires a tight regulation of immune responses and epithelial cell regeneration to combat environmental threats without the development of pathological inflammation. This is achieved by a highly adapted immune compartment, which is critically shaped by constant crosstalk with the microbiome, the key component of a balanced intestinal milieu (13). Ulcerative Colitis (UC) is a chronic or remitting inflammatory bowel disease (IBD) of the colonic mucosa affecting up to 0.42% of individuals in industrialized countries (14). Intestinal inflammation in UC patients typically spreads from the distal colorectum towards the proximal parts of the colon. Risk alleles for UC include genes important for epithelial barrier integrity or immune regulation, but explain only 7.5% of disease variants, while environmental factors and dysbiosis shape disease onset and severity (15). In fact, the pathophysiology of UC is driven by a triad of dysbiosis, barrier dysfunction and immune activation, but the actual sequence of patho-mechanistic events remains incompletely understood. In mice, UC can be mimicked by supplementation of the drinking water with dextran sodium sulfate (DSS). DSS disrupts the barrier layer, followed by entry of commensal bacteria into the mucosa and subsequent immune activation (16). Importantly, the DSS-sensitivity of mice highly depends on their hygiene status, again highlighting microbiome and host immune factors as critical determinants of disease severity (17). In addition, epithelial cell survival and regeneration are key parameters for barrier integrity, which ultimately reduces inflammation (18).

Based on the elevated FLNA editing in the colon (7) and the fact that specific mutations result in intestinal dysfunction (4), we hypothesized that the FLNA editing state may, similar to certain point mutations, impact intestinal development and/or immune homeostasis and specifically investigated the impact of genetically fixed FLNA editing states by assessing immunity, barrier function and microbiome composition in naïve- and DSS challenged FLNA mutant mice.

## Results

### FLNA editing impacts cell transcriptional profiles and inflammation in the healthy gut

The *Flna* mRNA is reported to be edited with 60 - 80% in the healthy murine colon (7, 11). Using FLNA mutant mice that exclusively express either edited FLNA^R^ or unedited FLNA^Q^ (8, 9), we first asked how the FLNA editing state impacts cellular transcriptomes under homeostatic conditions. We generated single cell suspensions derived from the epithelial layer (containing intraepithelial lymphocytes and enterocytes) and combined them with the lamina propria immune cell fraction from healthy FLNA^R^ and FLNA^Q^ colons and sorted viable CD45^-^ and CD45^+^ cells. Upon single-cell sequencing (Figure 1A), we identified nine clusters of epithelial cell (EC) states, including progenitors, transitional ECs (EC Trans I-III), distal ECs (Dist I-III) and proximal ECs (Prox I-II) as well as goblet cells, enteroendocrine cells and a few fibroblasts, tuft-, and smooth muscle cells (SMCs) (Figure 1B). The immune cell compartment consisted of different lymphocyte subsets including classical CD4+ T-helper cells, regulatory T- and Th17 helper cells, three subtypes of CD8+ T cells, a small cluster of innate lymphoid cells and B cells (Figure 1B). Myeloid cells were sparse, consisting of macrophages, monocytes, dendritic cells and a few neutrophils and mast cells (Figure 1B). Amongst structural cells, compositional analysis of ScSeq data suggested that edited FLNA^R^ colons harbored more transitional ECs, proximal ECs and goblet cells, but less differentiated distal ECs than unedited FLNA^Q^ colons (Figure 1C, left panel). Moreover, we isolated more atypical CD8+ T (γδT cells, Tinv) and CD4+ T helper cells from FLNA^R^ colons, at the cost of a slightly smaller B- and classical CD8+ T cell fraction (Figure 1C, right panel). However, flow cytometric analysis of CD45^+^ cells numbers did not reveal significant differences between groups with regards to CD8+ T cells (covering classical and atypical CD8+ T cells), CD4+ T, B cells or myeloid cells between the groups (Figure S1A and S1B). Differentially expressed genes (DEG) were determined for populations with at least ten cells per genotype, hence excluding fibroblasts, SMCs, tuft-, Th17- and all myeloid cells from analysis (Table S1). Profound differences were found in the EC compartment, namely in progenitors, early transitional EC, distal EC and goblet cells (Figure S1C-E). Interestingly, genes associated with epithelial barrier function (*Reg3b/g*, *Cldn2/8/15*, *Muc4*) (19–21) were upregulated in FLNA^R^ ECs, while FLNA^Q^ promoted a proinflammatory profile, with enhanced expression of genes involved in antigen presentation (e.g. *H2-Aa/Ab1/Eb1*, *B2m* (22)), interferon signaling (e.g. *Isg15*, *Stat1*, *Irf1/7/9*, *Bst2*, *Oas1a/2*, *Tap1)* (23) and neutrophil recruitment (*Cxcl1*) (24) (Figure 1D, S1C-E, Table S2). ECs also showed expression of genes involved in cell death. Gasdermins (*Gsdmc2/3/4*), caspase 1 (*casp1*), Asc (*pycard*), important pyroptosis related genes (25) and factors involved in ferroptosis, another type of inflammatory cell death, were highly expressed in FLNA^Q^ ECs (e.g. *Ncoa4*, *Fth1*) (26). Further, the FLNA editing state influenced genes associated with proliferation, cancer and stress response with some being upregulated in FLNA^R^ (e.g. *Naca*, *Myof*, *Smoc2*, *Hprt*) (27–30) and others being upregulated in FLNA^Q^ ECs, like the AP-1 transcription factor complex (*Atf3, Fos, Jun*) (31) (Figure 1D, S1B-D, Table S2). In contrast to structural cells, differences in immune cells were restricted to a few DEGs found in CD8+ T cells associated with inflammation and activation (*Tcrg, Trdc, Dnajb1, Hspa1a/1b*). Similar to ECs, we found *Hprt* upregulated in FLNA^R^ CD8+ T cells, while FLNA^Q^ immune cells expressed more *Fth1*, the ferritin heavy chain, suggesting a cell type independent impact of FLNA editing on *Hprt* and *Fth1* expression (Figure 1E, Table S2). Bulk-sequencing from distal colon homogenates confirmed *Hprt* upregulation in FLNA^R^ intestines (Table S3) and gene set enrichment analysis (32) confirmed a transcriptional signature that links FLNA^R^ with cell states found in proliferative disorders (DNA repair, Myc signaling) while suppressing inflammation (Figure 1F).

**Figure 1.**
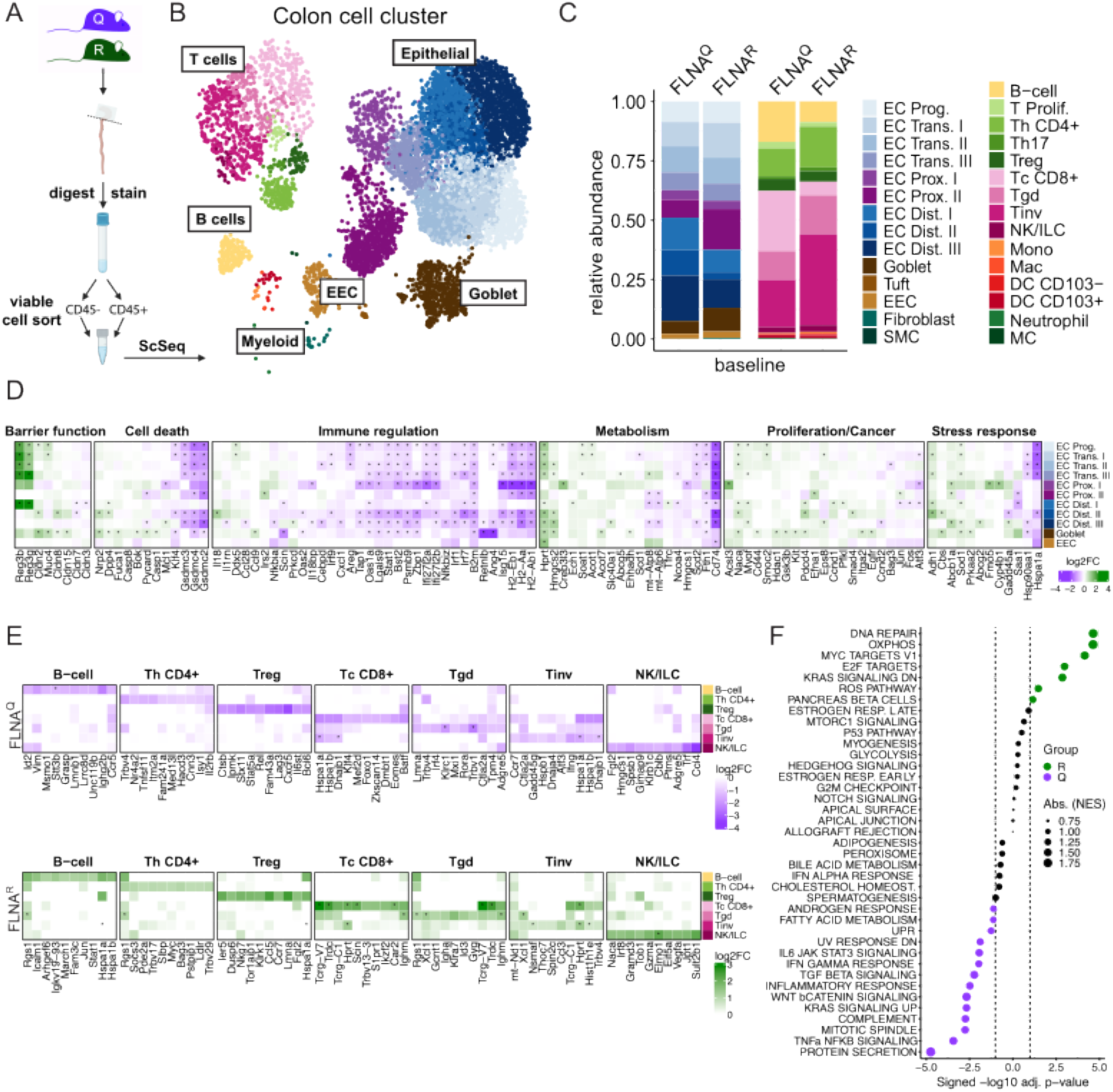
FLNA editing impacts cell transcriptional profiles and inflammation in the healthy gut. **A)** Scheme of scRNA-seq setup for FLNA^R^ and FLNA^Q^ mouse colons at basal conditions (3 mice pooled per group). **B)** Cell clustering of colon mucosal cells. **C)** Relative abundance of parenchymal (CD45-) and immune (CD45+) cells in FLNA^R^ and FLNA^Q^ intestines; left panel: epithelial cell progenitors (EC Prog), transitional ECs (EC Trans. I-III), proximal ECs (EC Prox. I-II), distal ECs (EC Dist. I-III), goblet cells, tuft cells, enteroendocrine cells (EEC), fibroblasts and smooth muscle cells (SMC); right panel: B-cells, proliferating T cells (T Prolif.), T helper CD4^+^ (Th CD4+), T helper 17 cells (Th17), regulatory T cells (Treg), classical cytotoxic CD8^+^ T cells (Tc CD8+), γδT cells (Tgd), invariant T cells (Tinv), natural killer/innate lymphoid cells (NK/ILC), monocytes (Mono), macrophages (Mac), CD103^-^ and CD103^+^ dendritic cells (DC), neutrophils and mast cells (MC) **D)** Heatmaps of transcriptomic analysis of DEGs in intestinal structural cells of FLNA^R^ and FLNA^Q^ mice grouped by cellular processes. Heatmap color represents the log2-fold increase of expression in FLNA^R^ (green) versus FLNA^Q^ (purple). Rows represent specific cell types. Asterisks represent adjusted values of *p*<0.05. **E)** Heatmaps of transcriptomic analysis of DEGs in intestinal immune cells of FLNA^R^ and FLNA^Q^ mice grouped by cell type. Heatmap color represents log2-fold change of expression in FLNA^R^ (green) and FLNA^Q^ (purple). Rows represent specific cell types. Asterisks represent adjusted values of *p*<0.05. **F)** Bubble plot representing top 40 MSigDB HALLMARK pathways enriched in bulk RNAseq transcriptomes of FLNA^R^ (green) and FLNA^Q^ (purple) cells. Circle sizes indicate the number of DEGs associated with the respective pathway.

Thus, FLNA editing state impacts gene expression in intestinal parenchymal- and immune cells. In ECs, edited FLNA^R^ promotes a transcriptome associated with proliferative cell states, while unedited FLNA^Q^ drives an inflammatory signature that may promote pyroptotic cell death.

### Edited FLNA^R^ is associated with an immune regulatory gut microbiome

Despite obvious differences in transcriptional states of intestinal cells, FLNA^R^ and FLNA^Q^ mutant animals appeared healthy and did not show signs of spontaneous colitis (8). Their bowel movements were normal, as indicated by an undistinguishable fecal pellet output in FLNA^Q^ and FLNA^R^ mice (Figure 2A-B) and no histological abnormalities (Figure 2C). The inflammatory state of intestinal cells depends on the integrity of the epithelial barrier, which shields the deeper immune-cell-rich tissue layers from the microbiota, but allows small metabolites to diffuse (13). However, we found no indication of epithelial barrier dysfunction in naïve animals. Comparable levels of FITC in the serum of both genotypes after oral gavage of FITC-dextran (Figure 2D) suggested a similar wall permeability. This was confirmed by e*x vivo* incubation of colons with FITC-dextran and quantification of the number of FITC+ cells present below the EC layer (Figure 2E-F).

**Figure 2.**
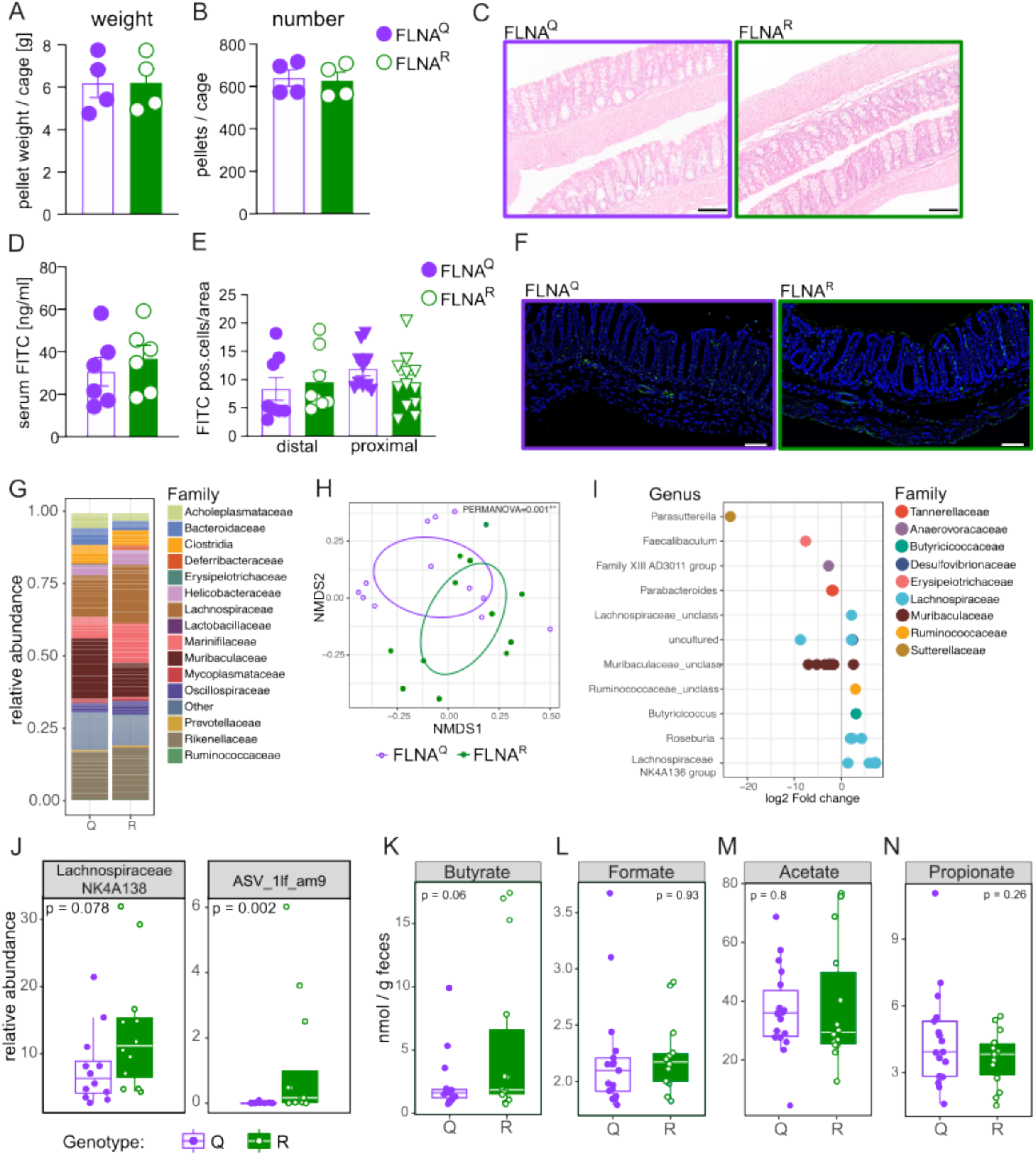
Edited FLNA^R^ is associated with an immune regulatory gut microbiome. **A)** Over-night fecal pellet output (weight) and **B)** average pellet number per cage (four animals per cage) of FLNA^Q^ and FLNA^R^ mice (*n*=16 per genotype, Student’s t-test). **C**) Representative H&E-stained colon sections from FLNA^Q^ and FLNA^R^ mice, scale bar=100µm. **D)** Serum fluorescence measurements after oral FITC-dextran gavage calculated as serum FITC in ng/ml (*n*=6, Student’s t-test). **E)** Number of FITC positive cells/area in cross sectioned colon tissue after *ex vivo* incubation of colons with FITC-dextran. Two proximal and two distal colon segments per mouse were analyzed (*n*=4, Student’s t-test). **F)** Representative images of colon cross-sections of distal colon. FITC+ cells in green, blue=DAPI, scale bar=100µm. **G)** Family-level relative abundance profiles of FLNA^R^ and FLNA^Q^ mice gut microbiomes. Families with relative abundances <2% are collapsed into the category “Other”. Each bar represents the average of multiple samples (*n*=6 per genotype). **H)** Ordination plot based on non-metric multidimensional scaling analysis (NMDS) of Bray-Curtis distances of microbiome profiles at the amplicon sequencing variant (ASV) level. Standard deviation ellipses and point dots representing each sample are depicted and are coloured by mouse genotype. Stress = 0.114. **I)** Differential abundant ASVs based on DESeq2 analysis (adjusted value of *p*<0.05, Wald test followed by Benjamini-Hochberg correction for multiple testing) between FLNA^R^ and FLNA^Q^ samples (log2-fold change > 2 denotes enrichment in FLNA^R^ mice; log2-fold change < −2 denotes enrichment in FLNA^Q^ mice). ASVs are assigned to genus (*y*-axis) and coloured by family. **J)** Relative abundance of the genus Lachnospiraceae NK4A136 group (left) and ASV_1lf_am9 (right) in fecal samples of FLNA^R^ (R) and FLNA^Q^ (Q) mice. **K-N)** Concentration of short-chain fatty acids butyrate, formate, acetate and propionate in fecal samples of FLNA^R^ and FLNA^Q^ mice measured by LC-MS (Student’s t-test). Error bars denote SD. Boxes represent the median, first and third quartile. Whiskers extend to the highest and lowest values that are within 1.5 times the interquartile range.

The intestinal epithelium and its immune system are fundamentally influenced by constant interaction with the microenvironment while the microbiome is reciprocally shaped by immune factors (33). Considering that in spite of an intact barrier, inflammatory signatures of intestinal epithelial cells were substantially reduced in FLNA^R^ animals, we hypothesized that FLNA mutant mice might harbor different microbiomes. We thus analyzed fecal pellets from FLNA^R^ and FLNA^Q^ mice, caged according to their genotypes after weaning, by 16S rRNA sequencing (Figure 2G). Indeed, the FLNA editing status influenced the microbiome, as its composition significantly clustered by genotype (Figure 2H). While both FLNA^Q^ and FLNA^R^ microbiomes were dominated by typical murine commensals (Figure 2G and 2I and 2J), fully edited FLNA^R^ mice showed a higher abundance of *Lachnospiraceae* and reduced *Parasutterella* and *Muribaculaceae* (Figure 2I and 2J). As *Lachnospiraceae* are known producers of short chain fatty acids (SCFA) with important immunomodulatory properties (34, 35), we measured SCFAs and indeed found higher butyrate levels in the feces of FLNA^R^ mice (Figure 2K), while propionate, acetate and formate were indistinguishable between genotypes (Figure 2L-N).

Taken together the FLNA editing state did not impact epithelial barrier integrity but was associated with a potentially more immunomodulatory microbiome in edited FLNA^R^ mice, which might be associated with the differential, less inflammatory transcriptional signatures we have observed in FLNA^R^ guts.

### FLNA editing protects mice from DSS-induced colitis

Considering the observed differences in transcriptional cell states and microbiomes, we next sought to test the impact of FLNA editing states on the susceptibility to colitis in an IL-10 deficient background. IL-10^−/−^ animals have an intact epithelial barrier, but progressively develop spontaneous colitis due to hyperinflammatory responses of macrophages to the microbiota, which are not counteracted by Treg derived IL-10 (36). Fixed FLNA editing states did not significantly impact spontaneous colitis upon IL-10 deficiency. By eight weeks of age, all IL-10^−/−^ animals already showed significant growth retardation and shortened colons independent of the FLNA genotype compared to wildtype animals (Figure S2A-B). No significant difference was seen upon severity scoring of intestinal tissues between FLNA^Q^ IL-10^−/−^ and FLNA^R^ IL-10^−/−^ mice, only FLNA^Q^ IL-10^−/−^ mice scored significantly worse than healthy controls (Figure S2C).

We next asked if a more severe disruption of intestinal homeostasis would unveil differences between FLNA mutants. To do so we chose the widely used DSS-induced colitis model (16). We treated FLNA^R^ and FLNA^Q^ mice with 2-2.5% of DSS in the drinking water for seven days, followed by three days of regular water (Figure 3A). Remarkably, we found that a complete lack of editing in FLNA^Q^ mice was associated with severe body weight loss, whereas constitutively edited FLNA^R^ animals were highly resistant to DSS (Figure 3B). The same pattern was seen for colon shortening, a reliable readout for intestinal inflammation, with pronounced shortening in FLNA^Q^ animals, whereas FLNA^R^ mice preserved their colon length (Figure 3C). Accordingly, FLNA^Q^ mice presented with a higher histopathological score than FLNA^R^ mice (Figure 3D-E) and this was most pronounced in the distal part of the colon, were colitis typically manifests (37) (Figure 3F-G). The pattern of FLNA^R^ protection and FLNA^Q^ sensitivity was observed for almost all scoring parameters, i.e., the degree of inflammatory cell infiltration, crypt damage, epithelial erosion and thickening (Figure 3H-K). Bulk RNA-sequencing of colon tissues from FLNA^R^ and FLNA^Q^ animals on day ten of DSS-induced colitis (at the peak of disease), confirmed that FLNA^R^ mice had less inflammation, as indicated by a reduced expression of inflammatory genes (*Il1b*, *Cxcl3*, *Cxcl2* or *Trem1*) (38). At the same time, FLNA^R^ mice further upregulated genes associated with cell differentiation, survival (e.g., Lgr5, Cbs, Ascl2) and barrier function (*Cldn2*) (Figure 3L-M and Table S4). Together, a fixed edited FLNA^R^ state renders mice highly resistant to DSS-induced colitis, while unedited FLNA^Q^ makes them more susceptible.

**Figure 3.**
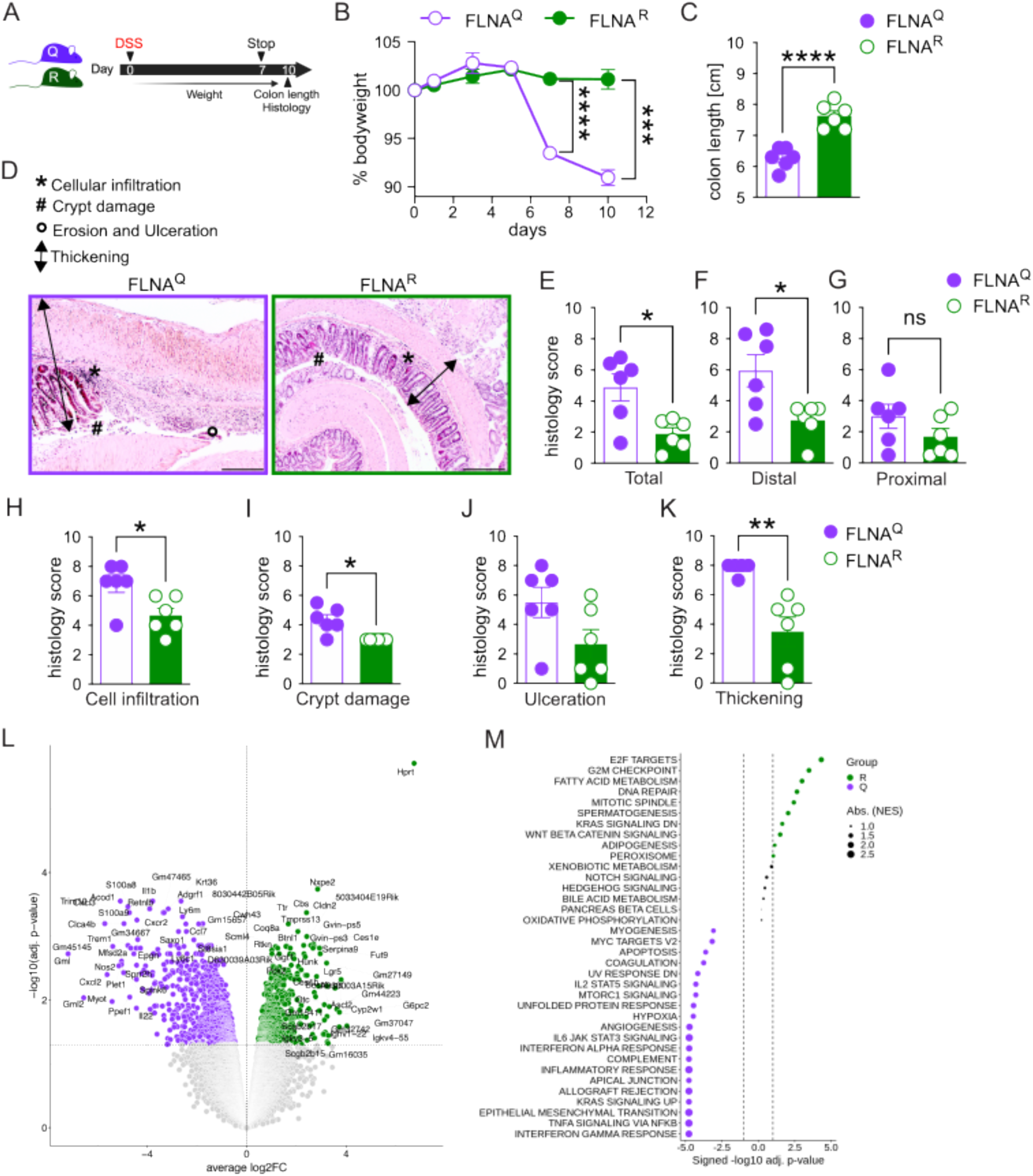
FLNA editing protects mice from DSS-induced colitis. **A)** Scheme of DSS-induced colitis experiment. **B)** Weight loss plotted in per cent of body weight compared to treatment start (2-way ANOVA, *F*=45.61, *DF*=5). **C)** Length measurements of colons at day ten (*n*=6, Student’s *t-*test). **D)** Representative images of histologic colon sections; bar = 500µm. **E)** Total **F)** Distal colon **G)** Proximal colon colitis histology score. A higher score means more severe colitis. (*n*=6, Mann Whitney U test). **H-K)** Individual colitis histology scoring parameters for the whole colon length: Cellular (immune cell) infiltration, crypt damage, erosion and ulceration of epithelium, thickening of colon wall; (*n*=6; Student’s *t-*test). **L)** Volcano plot (DEGs, padj ≤ 0.05) of FLNA^R^ versus FLNA^Q^ cells identified by DESeq2 analysis in whole distal colon bulk RNAseq. **M)** Bubble plot representing the top 40 MSigDB HALLMARK pathways enriched in bulk transcriptomes of FLNA^R^ (green) and FLNA^Q^ (purple) mice at day ten of DSS challenge. Circle sizes indicate the number of DEGs associated with the respective pathway. ****p<0.0001; ***p<0.001; **p<0.01; *p<0.05. Error bars show SEM.

### FLNA^R^ reduces early inflammatory cytokine production and intestinal neutrophil accumulation

We next wanted to gain mechanistic insights into the differential DSS susceptibility of FLNA^Q^ and FLNA^R^ mice. Analogous to what we did before in healthy animals, we analysed sc-RNAseq of viable CD45^+^ and CD45^-^ cells on day five of DSS-induced colitis, when mice are not severely sick yet, but colon shortening already manifests in FLNA^Q^ animals (Figure S2D). Cells were clustered as before (Figure S2E). An enrichment of fibroblasts and granulocytes and a reduction in goblet- and B cells was apparent in both genotypes upon DSS challenge (Figure 1C and 4A). In FLNA^R^ animals we observed a higher frequency of mature EC, which may indicate reduced cell loss, and a lower neutrophil abundance (Figure 4A). DEGs were determined for all populations except SMC, tuft- and proliferating T cells (Table S5).

**Figure 4.**
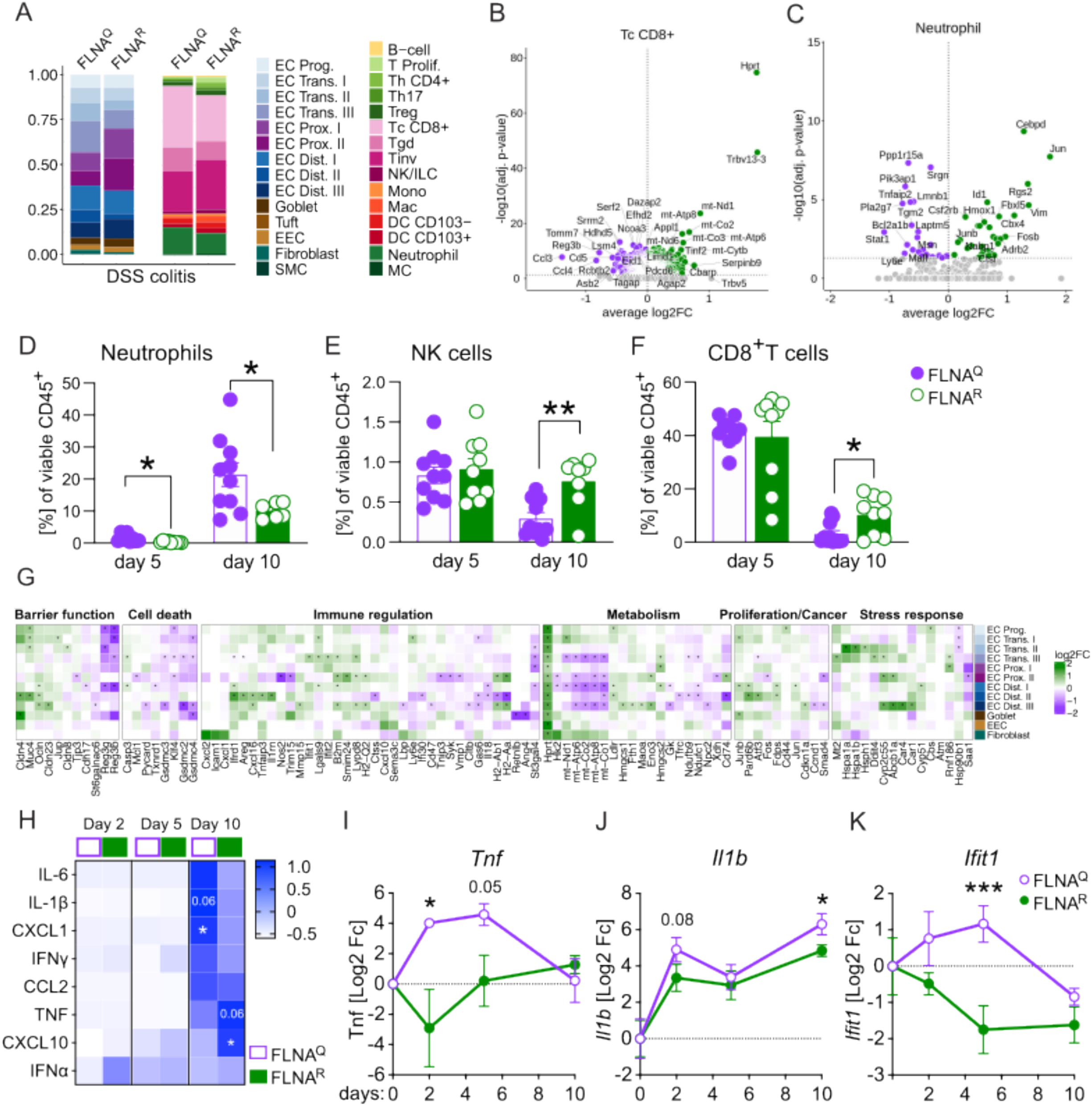
FLNA^R^ reduces early inflammatory cytokine production and intestinal neutrophil accumulation. **A)** Relative numbers of parenchymal (CD45-) and immune (CD45+) cells in FLNA^R^ and FLNA^Q^ intestines; left panel: epithelial cell progenitors (EC Prog), transitional ECs (EC Trans. I-III), proximal ECs (EC Prox. I-II), distal ECs (EC Dist. I-III), goblet cells, tuft cells, enteroendocrine cells (EEC), fibroblasts and smooth muscle cells (SMC); right panel: B-cells, proliferating T cells (T Prolif.), T helper CD4^+^ (Th CD4+), regulatory T cells (Treg), T helper 17 cells (Th17), classical cytotoxic CD8^+^ T cells (Tc CD8+), γδT cells (Tgd), invariant T cells (Tinv), natural killer/innate lymphoid cells (NK/ILC), monocytes (Mono), macrophages (Mac), CD103^-^ and CD103^+^ dendritic cells (DC), neutrophils and mast cells (MC). **B-C)** Volcano plots (DEGs, p-adj. ≤ 0.05) of FLNA^R^ versus FLNA^Q^ cells identified by DESeq2 analysis for classical CD8^+^ T cells and neutrophils. **D-F)** Relative numbers of intestinal neutrophils, NK cells and CD8^+^ T cells (Student’s *t-*test). **G)** Heatmaps of selected DEGs in intestinal epithelial cells of FLNA^R^ and FLNA^Q^ mice after five days of DSS grouped by cellular processes. Heatmap color represents log2-fold increase of expression in FLNA^R^ (green) and FLNA^Q^ (purple). Rows represent cell types of epithelial cells. Asterisks represent significant p-adj.<0.05 values of log2-fc. **H)** Heatmap of cytokine levels (pg/ml) in mouse colon homogenates shown as z-score across both groups and timepoints (2-way ANOVA and Šídák multiple comparisons test). **I-K)** Gene expression analysis by real-time qPCR of FLNA^R^ and FLNA^Q^ colon homogenates. CT values relative to *Gapdh* (2^-ddCT), shown as log2fold change compared to day zero per timepoint (2-way ANOVA and Tukey’s multiple comparisons test). ****p<0.0001; ***p<0.001; **p<0.01; *p<0.05. Error bars show SEM.

In contrast to naïve mice, FLNA^R^ and FLNA^Q^ immune cells showed substantial differences during colitis with most DEGs found in classical cytotoxic CD8+ T cells (Tc CD8+) (Table S6 and Figure 4B). While FLNA^R^ Tc CD8+ cells expressed enhanced levels of T cell receptor (TCR) components (*Trbv13-3, Trbv5*), FLNA^Q^ Tc CD8+ cells upregulated factors associated with TCR regulation (e.g. *Cd5*) (39), recruitment (*Ccl3, Ccl4*) and activation (*Reg3b*) (19, 40) (Figure 4B). While we also found a substantial number of DEGs in the other two CD8+ T cell subsets, other immune cell types were largely unaffected by the FLNA editing status, with one exception: FLNA^R^ neutrophils exhibited a transcriptional profile associated with anti-inflammatory properties (e.g. *Adrb2, Hmox1, Rgs2)* (41), cell survival (e.g. *Jun, Junb, Fos*) and differentiation (e.g. *Id1*, *Csf2rb*) (42). FLNA^Q^ neutrophils on the contrary upregulated inflammatory genes (e.g. *Stat1, Ly6e, Tnfaip2*) (43) (Table S6 and Figure 4C). To confirm potential compositional differences and to assess the dynamics of immune cell infiltration, we analyzed intestinal cell infiltration over time during DSS-induced colitis. While on day two we did not detect increased leukocyte numbers compared to healthy mice, infiltration started to rise by day five – a timepoint when also a trend towards increased total leukocyte numbers was observed in FLNA^Q^ animals - and got even higher by day ten (Figure S2F). By day two the cell composition was almost indistinguishable from healthy mice, except for a small, yet significant increase in monocytes of FLNA^R^ intestines (Figure S1A-B and S2G). This difference was not visible any longer by day five of DSS treatment when, in accordance to the ScSeq data, neutrophil infiltration was significantly lower in protected FLNA^R^ than FLNA^Q^ colons and this trend continued until day ten (Figure 4D and S2G). While FLNA^R^ mice significantly preserved their intestinal lymphocyte pools, a substantial depletion of NK and CD8+ T cells was observed in FLNA^Q^ mice (Figure 4E-4F). Other immune cell abundances such as B cells, T helper cells, macrophages and DCs were unaffected (Figure S2H-S2J).

Under homeostatic conditions, FLNA^R^ structural cells expressed more genes associated with barrier function (*Reg3b/g*, *Cldn2/8/15*, *Muc4*) (Figure 1). During colitis a higher expression of *Muc4* was maintained and in addition *Cldn4*, *Ocln* and *Jup*, important tight- and adherens junction components, were upregulated (44). *Reg3g*/*3b* were now higher expressed in FLNA^Q^ enterocytes, which may reflect an increased stimulation of the epithelium by the microbiota (Figure 4D) (19). While gasdermin transcription remained higher in FLNA^Q^ cells, some FLNA^R^ ECs showed elevated caspase 3 transcription, suggesting that FLNA^R^ rather promoted a transcriptional state associated with apoptosis, while FLNA^Q^ promoted pyroptotic cell death (25). Again, the FLNA editing state influenced the expression of proliferation-cancer- and stress response associated genes (*Junb*, *Jun*, *Pard6b*, *Atf3, Hspa1a* and -*b, Cyp2c55*, *Abcb1a)* (45, 46) (Figure 4G). We then assessed protein and mRNA levels of inflammatory cytokines in colon homogenates. Protein cytokine levels were still low on day two and five but increased by day ten in both genotypes. Concentrations of IL-1β and the neutrophil chemoattractant CXCL1 were elevated in susceptible FLNA^Q^ mice, while protected FLNA^R^ colon homogenates contained more TNF and the IFN-inducible factor CXCL10 (Figure 4H). On RNA level, distinct inflammatory patterns were detectable already as early as day two, with reduced expression of *Tnf, Il1b* and *Ifit1* in FLNA^R^ compared to FLNA^Q^ mice (Figure 4I-4K).

Together, we conclude that the FLNA editing state modulates early inflammatory responses to DSS, thus influencing the intestinal cytokine milieu and immune cell infiltration. Considering the importance of neutrophils in driving DSS-induced tissue damage (47) and the finding of transcriptional and infiltration differences of neutrophils, we speculated that differential recruitment and functional state of neutrophils contributed to the observed protection in fully edited FLNA^R^ animals.

### A potentially protective microbiome does not explain DSS-resistance of FLNA^R^ mice

SCFAs and butyrate in particular have been associated with anti-inflammatory effects in IBD and mouse models of intestinal inflammation (34). Thus, it was tempting to speculate that the butyrate-enriched microbiome of FLNA^R^ animals may contribute to their DSS-resistance (Figure 2). When we tested, if microbiota and butyrate content were associated with the phenotypic outcome of the DSS treatment in FLNA mutant mice, it was not surprising to find a relationship between the community composition and fecal butyrate levels (Figure 5A). However, the community composition in naïve mice strikingly also predicted the colitis score in FLNA^R^ and FLNA^Q^ mice upon subsequent DSS treatment (Figure 5A). Among all genera, the *Lachnospiraceae NK4A136* group was strongly associated with butyrate levels (Figure 5A), likely because organisms from this genus are closely related to several butyrate-producing microbes (48). We further identified an ASV within this genus (ASV_1lf_am9, 96% identical to *Lachnoclostridium pacaense*) whose abundance positively correlated with butyrate levels (Figure S3A) and negatively with the colitis score (Figure S3B). As these results suggested a connection between microbiome and colitis severity, we next experimentally tested a potential causality of the microbiome in the DSS-resistance of FLNA^R^ mice. First, we depleted the microbiota of the mice for three weeks with a cocktail of antibiotics (Figure 5B), reflected by a reduction in the total number of bacteria (Figure S3C-S3D), and a drop in alpha diversity (Figure S3E). However, when microbiome-depleted- and control mice were then challenged with DSS (Figure 5B), FLNA^R^ animals were still significantly and similarly protected from body weight loss and colon shortening (Figure 5C-5D). In a second approach, we co-housed FLNA^Q^ and FLNA^R^ to align their microbiomes prior to DSS challenge (Figure 5E). However, cohousing also did not abrogate the difference in colitis severity between FLNA^Q^ and FLNA^R^ animals, as FLNA^R^ mice still lost less body weight and preserved their colon lengths (Figure 5F-5G). Thus, the potentially protective microbiome, we had observed in FLNA^R^ animals, can only contribute to, but is not causal for their protection from DSS-induced colitis.

**Figure 5.**
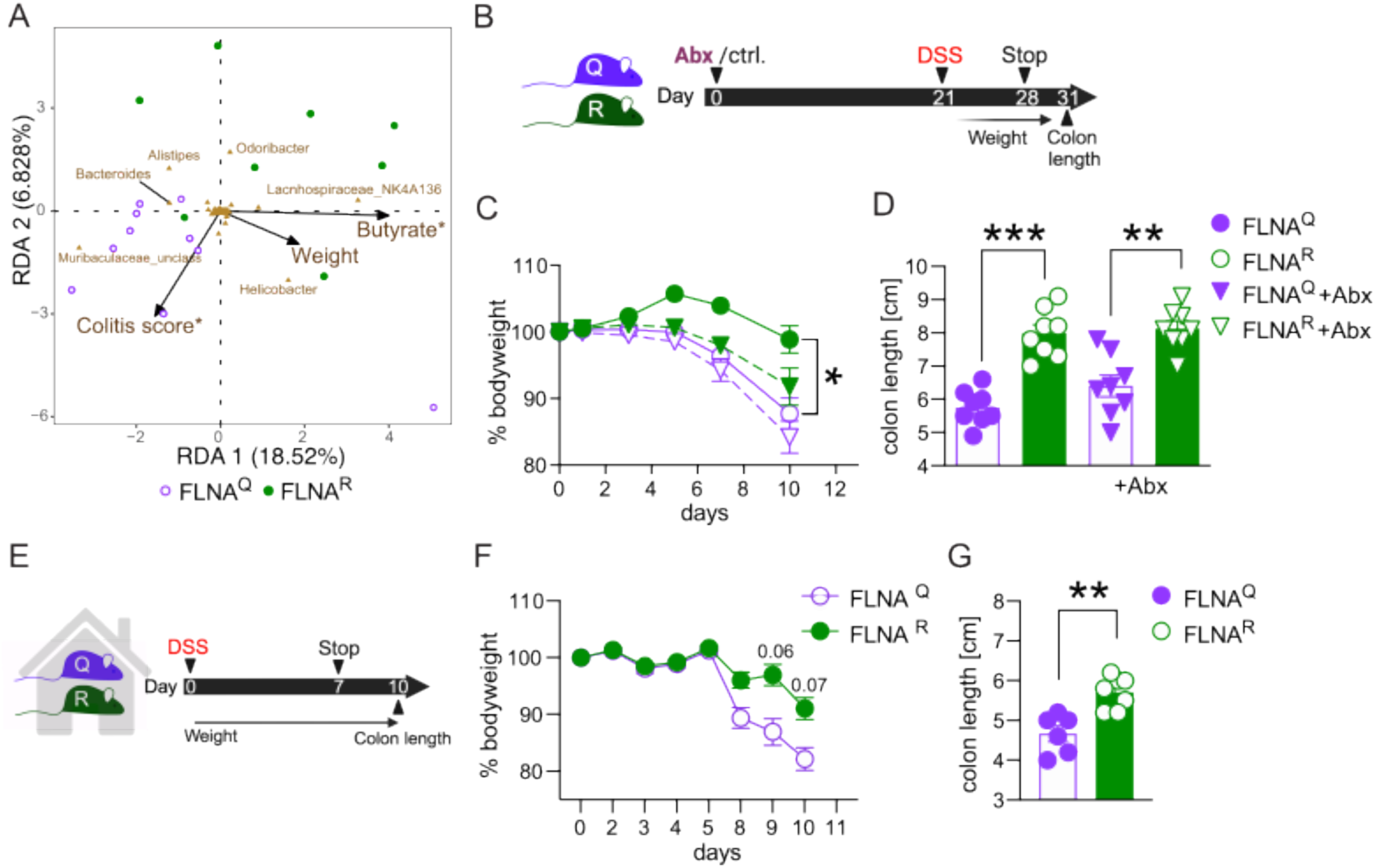
A potentially protective microbiome does not explain DSS-resistance of FLNA^R^ mice. **A)** Outcome predictive redundancy analysis (RDA) of the prokaryotic community of FLNA^Q^ (purple dots) and FLNA^R^ (green dots) mice. The relation of the community composition (at the genus-level) with the environmental variables: butyrate, colitis score and body weight loss (% of initial body weight) is shown. Direction and length of arrows shows the correlational strength between the abundance of each prokaryotic genus and the environmental variable. Asterisks denote environmental factors that are statistically significant (PERMANOVA; *p*<0.05). **B)** Scheme of antibiotic (Abx) depletion and DSS challenge. **C)** Weight loss plotted in % of bodyweight compared to treatment start for FLNA^R^ and FLNA^Q^ mice with (dashed line) or without (solid line) microbiome depletion. **D)** Length measurements of colons at day ten (1-way ANOVA, *F*=21.20, *DF*=28). **E)** Scheme of cohousing and DSS treatment. **F)** Weight loss curves for cohoused FLNA^R^ and FLNA^Q^ mice (2-way ANOVA, *F*=2.941, *DF*=5). **G)** Length measurements of colons for cohoused FLNA^R^ and FLNA^Q^ mice at day ten (Student’s t-test). ****p<0.0001; ***p<0.001; **p<0.01. Error bars show SEM.

### Edited FLNA^R^ in myeloid cells protects from colitis

Having excluded a major contribution of the microbiome to the disease phenotype, we next sought to dissect which cell types mediate the differential DSS susceptibility upon fixation of the FLNA editing state. Higher levels of FLNA editing (65-100%) have earlier been described in mucosal fibroblasts and endothelium and low-medium levels (16-50%) in smooth muscle cells and keratinocytes, while immune cells and epithelial cells mainly express unedited FLNA under homeostatic conditions (12).

Considering the high levels of endothelial editing and our finding of a significant reduction in intestinal neutrophil accumulation of DSS resistant FLNA^R^ mice, we next hypothesized that FLNA editing may impact inflammation-induced leukocyte trafficking via its influence on endothelial cells and/or hematopoietic cells. To investigate this, we made use of a newly generated mouse, which constitutively expresses unedited FLNA^Q^ but induces edited FLNA^R^ upon Cre-recombination (FLNA^QiR^) (Figure 6A). We first generated FLNA^QiR^ Vav1Cre^+/−^ mice to specifically express FLNA^R^ in hematopoietic- and endothelial cells (49) and verified FLNA^R^ expression in the spleen and unedited FLNA^Q^ in tissues with small immune cell populations (intestine, stomach) (Figure S4A). When FLNA^QiR^ VavCre^+/−^ and FLNA^QiR^ control animals were subjected to the DSS regime, we found that FLNA^QiR^ VavCre^+/−^ mice were protected, indicated by significantly preserved colon length and an improved colitis score as compared to unedited FLNA^QiR^ animals (Figure 6B-C). Although a similar trend was visible for the bodyweight loss, these differences did not reach statistical significance (Figure S4B).

**Figure 6.**
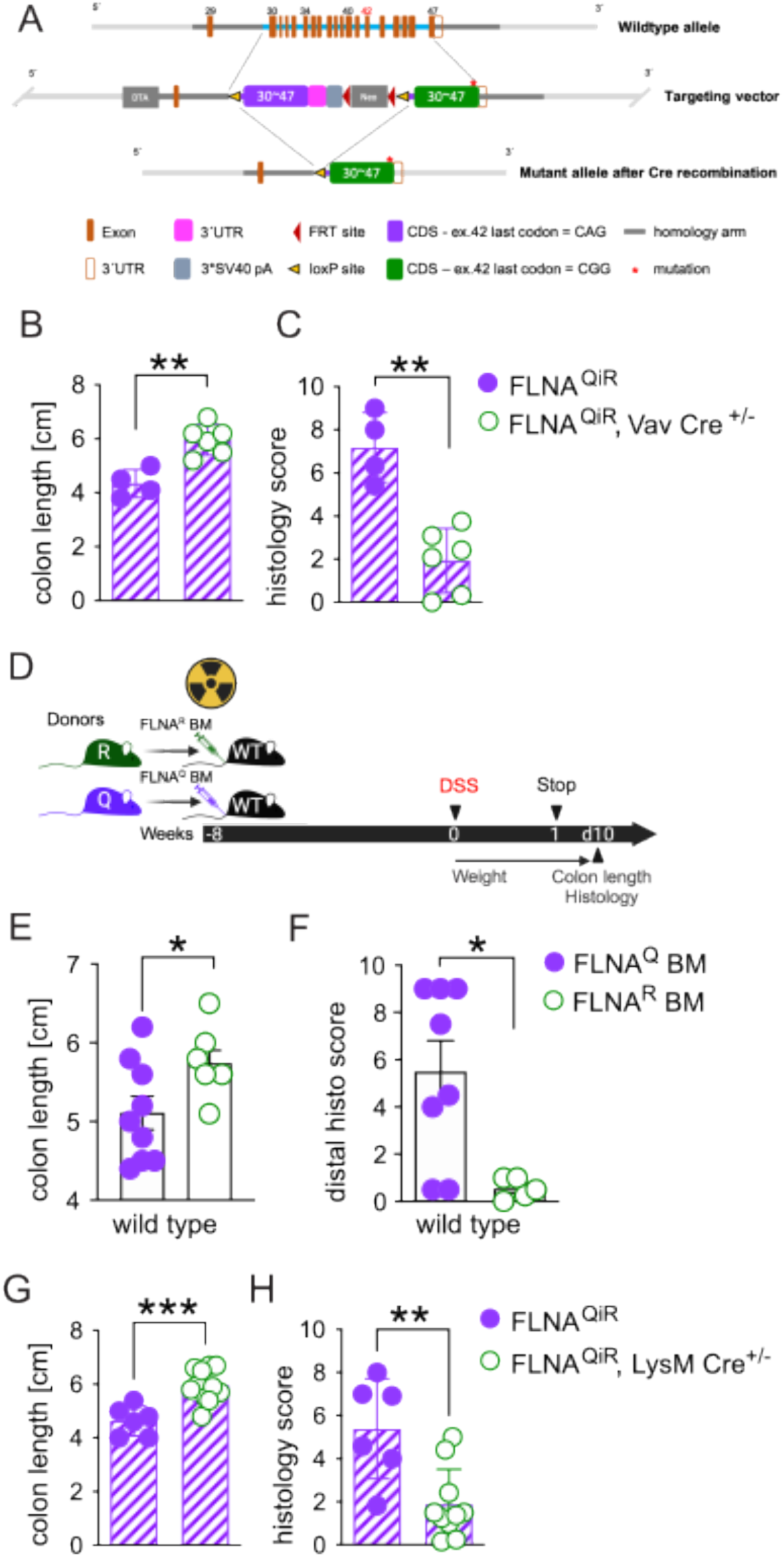
Edited FLNA^R^ in myeloid cells protects from colitis. **A)** Organization of the FLNA WT allele, the targeting vector, and the inducible FLNA^QiR^ -allele. The targeting vector cassette replaced exons 30-47 by homologous recombination. The loxP flanked unedited CDS (purple) & the Neo-cassette are then deleted by Cre-recombinase. **B-C)** Colon length measurements and total histology colitis scores of FLNA^QiR^ and FLNA^QiR^ Vav Cre^+/−^ mice at day ten of DSS challenge (*n=*4 for QiR and *n=*6 for QiR-Cre, Student’s *t-*test and Mann Whitney U test). **D)** Scheme of BM transplant experiments followed by DSS treatment. **E-F)** Colon length measurements and colitis histology score of lethally irradiated wild type mice reconstituted with either FLNA^Q^ or FLNA^R^ BM (*n=*9 for Q and *n=*6 for R, Student’s *t-*test and Mann Whitney U test). **G-H)** Colon length measurements and total histology colitis score of FLNA^QiR^ and FLNA^QiR^ LysM Cre^+/−^ mice at day ten of DSS challenge (*n=*6 for QiR and *n=*10 for QiR-Cre, Student’s *t-*test and Mann-Whitney U test). ***p<0.001; **p<0.01; *p<0.05. Error bars show SEM.

To narrow down the effects of a fixed FLNA editing state to endothelial or hematopoietic cells, we next made use of a bone marrow transplant approach. To assess potential impacts of the FLNA editing state on hematopoietic cell homing and bone marrow reconstitution, we first analyzed repopulation efficiencies in a setup where we transplanted Ub-GFP (which resembles wild type), FLNA^Q^ or FLNA^R^ bone marrow into lethally irradiated WT animals. Blood and bone marrow leukocyte counts were indistinguishable between groups and we found no signs of inflammation (Figure S4C-D), showing that all genotypes had repopulated similarly by 8 weeks post transplantation. As expected, T cells were replaced with the lowest efficiency (around 75-80%), which is due to the radioresistant properties of certain T cell subsets (50). Most efficient replacement was observed for neutrophils and B cells (Figure S4E-F). We thus concluded that the FLNA editing state did not influence bone marrow reconstitution after irradiation and next transplanted bone marrow from FLNA^Q^ and FLNA^R^ mice into WT mice followed by DSS challenge (Figure 6D). Although we did not observe any difference in bodyweight loss (Figure S4G), transplantation of FLNA^R^ bone marrow still induced protection as indicated by a preserved colon length (Figure 6E) and improved colitis scores, particularly in the distal colon (Figure 6F and S4H). Thus, these data surprisingly revealed that hematopoietic FLNA^R^ is sufficient to protect from colitis.

Based on this result and the notion that neutrophils may drive potential protective effects - FLNA^R^ neutrophils exhibited an anti-inflammatory transcriptional profile and reduced migration to the tissue during colitis (Figure 4C and 4E) - we crossed FLNA^QiR^ and LysMCre animals. Doing so we generated mice that either lack editing or express constitutively edited FLNA^R^ only in myeloid cells, including neutrophils (51) and macrophages. Strikingly, FLNA^QiR^ LysMCre^+/−^ mice recapitulated the phenotype we had observed in FLNA^QiR^ VavCre^+/−^ animals and irradiated mice that were reconstituted with FLNA^R^ bone marrow. FLNA^QiR^ LysMCre^+/−^ significantly preserved colon length (Figure 6G) and tissue integrity (Figure 6H), and additionally, lost less bodyweight (Figure S4I). Importantly, a depletion of CD8^+^ T cells using an established depletion regime with an anti-CD8 antibody (52) (Figure S4J) did not abrogate the differences in colon length between FLNA^Q^ and FLNA^R^ animals (Figure S4K-M). We thus concluded that fixed FLNA^R^ in myeloid cells reduces inflammatory cytokines and neutrophil tissue infiltration during DSS challenge, which drives protection from severe colitis.

### A fixed fully edited FLNA^R^ in myeloid cells shifts cellular properties and could be therapeutically exploited in colitis

Having established that a fixed FLNA^R^ state in myeloid cells, is protective during DSS-induced colitis, we wanted to follow the dynamics of FLNA editing levels during colitis in inflamed wild type colon tissue. Using *Flna* editing site amplicon sequencing, we first assessed FLNA editing in distal colon homogenates in homeostasis as well as on day two, five and ten of acute colitis. FLNA was edited to 60% at baseline and this was maintained on day two post DSS challenge.

From day five on, the FLNA editing ratio decreased until day ten, where we only found editing of 30% (Figure 7A) and this went along with a reduction of *Flna* expression (Figure 7B upper panel and S5A). Reduced editing and *Flna* expression levels were associated with an increased expression of classical proinflammatory cytokines and chemokines like Il1b, Cxcl1, or Il6, which are typically induced during colitis (Figure 7B lower panel, S5B-D). Expression levels of *Adarb1* - encoding for ADAR2 that catalyzes *Flna* mRNA A-to-I deamination (7) - did neither correlate with *Flna* expression nor editing levels and remained low but stably expressed during the course of colitis (Figure 7B lower panel and S5F). Only a trend for decreased Adar (ADAR1) expression, was observed, but did not correlate with *Flna* editing levels (Figure 7B lower panel, S5E). Together, these data suggested that the downregulation of FLNA editing during colitis was not caused directly by a reduction of the editase (ADAR2), but rather associated with inflammation and, most likely, tissue accumulation of unedited immune cells, and/or loss of highly edited cells.

**Figure 7.**
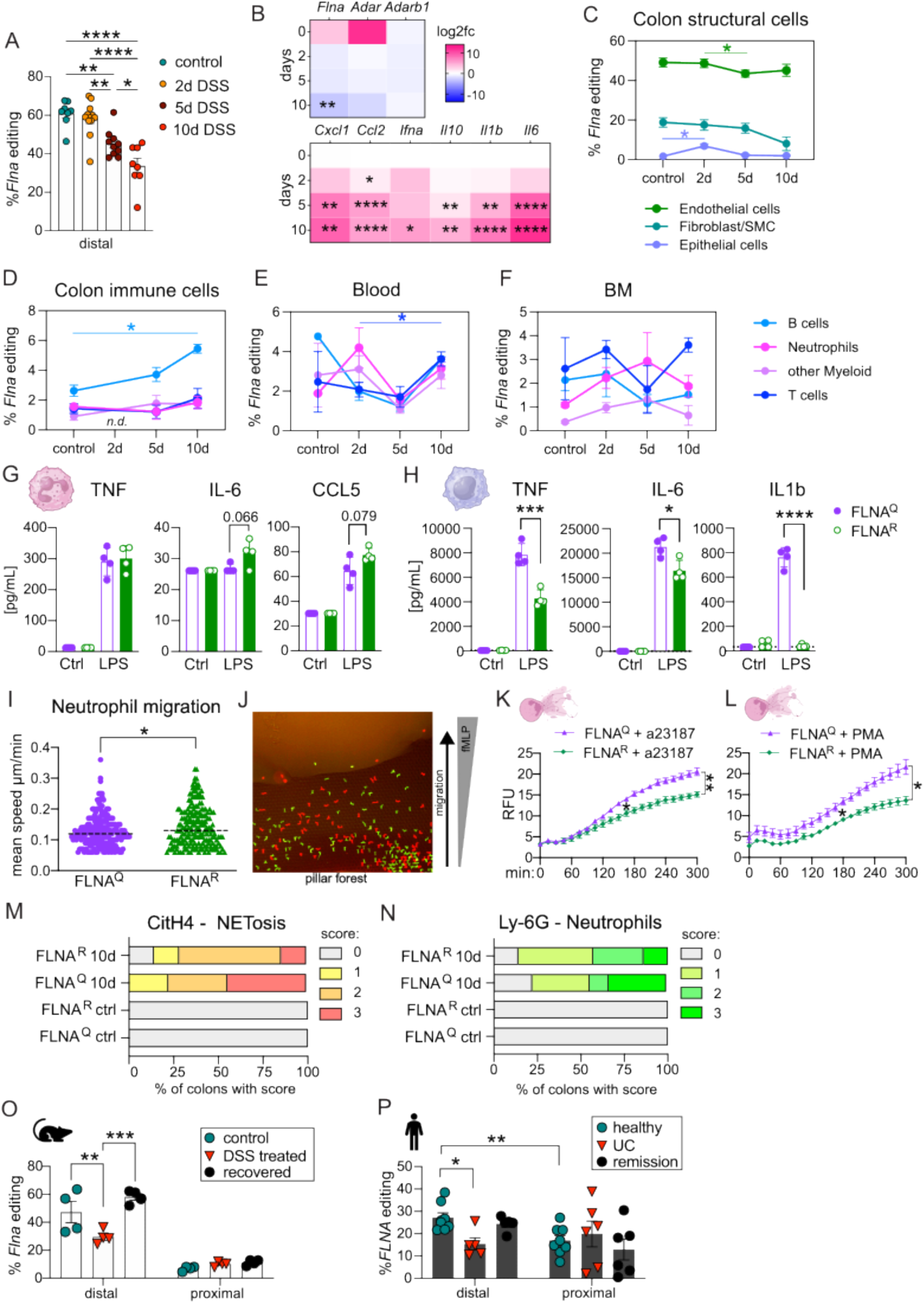
A fixed fully edited FLNA^R^ in myeloid cells shifts cellular properties and could be therapeutically exploited in colitis. **A)** *Flna* mRNA editing status of whole distal colon of wildtype mice determined by RT-PCR amplicon Illumina sequencing on days zero, two, five and ten of DSS challenge (1-way ANOVA, *F=*19.09, *DF*=3). **B)** Gene expression analysis by real-time qPCR from colon homogenates. CT values relative to *Gapdh* (2^-ddCT), shown as log2fold change compared to day zero depicted as heatmap at different timepoints of DSS treatment (Mixed-effects analysis followed by Tukey’s multiple comparisons test). **C-F)** *Flna* mRNA editing status of sorted cell populations of two-six pooled wildtype mouse colons, BM and blood determined by RT-PCR amplicon Illumina sequencing on days zero, two, five and ten of DSS challenge (Mixed-effects analysis followed by Tukey’s multiple comparisons test). **G-H)** Bar graphs of cytokine levels (pg/ml) in supernatants from LPS stimulated mouse neutrophils and BMDMs (Student’s *t-*test). **I)** Neutrophil migration speed in a pillar forest of a microfabricated PDMS device towards a fMLP gradient (Student’s *t-*test). **J)** Exemplary image of CSFE- and TAMRA-stained FLNA^Q^ and FLNA^R^ neutrophils within the device. **K-L)** Quantification of DNA release as relative fluorescence units to measure NETosis of isolated neutrophils post stimulation with a23187 or phorbol 12-myristate 13-acetate (PMA) over time. Time course data were plotted in a line graph with mean ± SEM (2-way ANOVA followed by Tukey’s multiple comparisons test, *F*=11.94, *DF*=60). **M-N)** Scoring of histological colon sections stained for NETs by anti-citrullinated histone 4 (citH4) or neutrophils by anti-mouse Ly-6G. **O)** *Flna* editing levels in intestinal tissues of wildtype mice: healthy mice, mice with acute colitis (day ten of DSS) or mice recovered from colitis (day 21) were analysed for *Flna* editing levels in distal and proximal colon separately (*n=*4; 1-way ANOVA, *F*=9.208, *DF*=18). **L)** *FLNA* editing levels in human colon biopsies of healthy donors (*n=*9), patients suffering from active UC (*n=*6) and patients in remission from active UC (*n=*6). Distal and proximal colon biopsies were analysed separately (1-way ANOVA, distal: *F*=7.326, *DF*=15); comparison healthy-distal vs healthy-proximal (Student’s t-test). ****p<0.0001; ***p<0.001; **p<0.01; *p<0.05. Error bars show SEM.

To address, which cells edited FLNA under homeostatic conditions and during colitis, we took the challenge to sort different structural- and immune cell populations from colons of healthy- and DSS challenged animals. As seen in our scSeq datasets, our initial colon preparation protocol was optimized for the isolation of intraepithelial lymphocytes, lamina propria immune- and intestinal epithelial cells (53), but failed to obtain viable intestinal endothelial cells. We therefore performed separate experiments from which we isolated CD31+ intestinal endothelial cells and a CD45-EpCam-CD31-cell fraction that is enriched for fibroblasts and SMCs. Epithelial cells expressed only low levels of edited FLNA^R^, but higher levels (up to 50%) were found in endothelial- and moderate levels in the Fibroblast/SMC fraction (up to 20%). Endothelial cells and Fibroblast/SMCs reduced their editing frequencies during colitis, although changes for Fibroblast/SMCs did not reach statistical significance (Figure 7C). Analysis of intestinal neutrophils on day two post DSS challenge was hampered by insufficient cell numbers retrieved (even upon pooling of multiple colons). We thus alternatively sorted immune cells from blood and bone marrow. Editing was generally low, but not absent, in all isolated intestinal immune cell populations, namely T cells, B cells, neutrophils and a fraction enriched for CD11b+ myeloid cells other than neutrophils. Except for a significant elevation of FLNA editing in B cells by day ten of DSS colitis, editing fluctuated between 0-5% and was not affected by DSS treatment (Figure 7D). A similar result was found in blood and bone marrow (Figure 7E and 7F). Accordingly, and in contrast to ubiquitously expressed *Adar*, we only found high levels of *Adarb1* in fibroblasts and some few enterocytes, while it was not detectable in most immune cells in FLNA^Q/R^ colons and in a ScSeq dataset derived from WT animals (Figure S5G and S5H). Together, these data showed that a fixed FLNA^R^ state in neutrophils and other myeloid cells conferred protection from colitis, despite a low FLNA editing ratio in these cells, whereas, similar to other tissues (12), FLNA is mainly edited in colon endothelium, fibroblasts and SMCs. We thus propose that targeted myeloid cell FLNA editing reveals a new option of therapeutic intervention for intestinal inflammatory diseases.

To better characterize the consequences of an artificial FLNA^R^ state in myeloid cells, we assessed the functional impact of FLNA^R^ on critical myeloid cell effector functions using primary bone marrow neutrophils and bone marrow derived macrophages (BMDM) isolated from either FLNA^Q^ or FLNA^R^ mice. Upon stimulation with the Toll-like receptor 4 ligand, Lipopolysaccharide (LPS), neutrophils of both genotypes produced even amounts of TNF, but FLNA^R^ neutrophils showed a trend to release more of the inflammatory cytokine IL-6, and the chemokine CCL5 (Figure 7G). In BMDMs, which are much more potent cytokine producers than neutrophils, we interestingly found the opposite effect. FLNA^R^ BMDMs were significantly hampered in TNF, IL-1β and IL-6 release in response to LPS as compared to FLNA^Q^ BMDMs (Figure 7H), showing that even within the myeloid cell compartment, FLNA editing exerts highly cell-type specific effects. Using microfluidic chambers, we assessed neutrophil migration towards an fMLP gradient. While both editing states allowed for directed migration of neutrophils, FLNA^R^ neutrophils migrated slightly, yet significantly, faster than FLNA^Q^ neutrophils. However, this difference was only measurable during the first hour of observation (Figure 7I and 7J), before extreme collective dynamics (swarming behavior) masked any cell-autonomous differences. It remains elusive whether such subtle differences between the genotypes could impact physiology. The release of neutrophil extracellular traps (NETs), the controlled expulsion of extracellular fibers containing nuclear DNA, histones and antibacterial granule components, is a key feature of activated neutrophils in inflammatory settings (54). During acute infection, NETosis can hinder pathogen dissemination, but also promotes immunothrombosis, inflammation, barrier- and tissue damage (55). In return, inflammatory mediators such as IL-1ß and TNF have been shown to enhance NET release (56, 57). While we had observed a rather enhanced activity of FLNA^R^ neutrophils regarding cytokine release and migration, we strikingly found that FLNA^R^ neutrophils were less prone to release NETs in response to a23187 and PMA, two established NET-inducers (58) (Figure 7K and 7L). Accordingly, we also found reduced NET formation *in vivo* during colitis, suggesting that a fixed FLNA^R^ state in myeloid cells may promote a micromilieu that contains inflammation and NETosis (Figure 7M-N and S5I).

To foster the mechanistic interpretation of our findings, we investigated potential differences in cell-cell interaction using ChellChat analysis based on our single cell datasets (59). This revealed similar signaling patterns (with regards to the inferred signaling pathways), but distinct interaction strengths between FLNA^Q^ and FLNA^R^ cell populations. In particular, the interaction of the intracellular adhesion molecule-1 (ICAM1) with different integrins seemed affected by the FLNA editing state. While the differences are rather subtle, they were most pronounced within the myeloid cell compartment with FLNA^Q^ neutrophils showing a stronger autocrine signaling and an increased interaction with DCs and MonoMacs (a mixed population of monocytes and macrophages). In contrast, FLNA^Q^ fibroblasts signaling to neutrophils via the ICAM1-integrin axis was reduced (Figure S5J). FLNA^Q^ and FLNA^R^ MonoMacs differed in their signaling through semaphorin 4D (Sema4D/CD100), an immunoregulatory transmembrane protein signaling to Cd72 or plexins in immune and non-immune cells, respectively (60). The interaction of macrophage Sema4d and Cd72 on other immune cells was more pronounced for FLNA^R^ cells. Similar to neutrophils, ICAM1-integrin signaling was slightly enhanced from FLNA^Q^ immune cells to MonoMacs, while reduced from FLNA^Q^ fibroblasts to MonoMacs (Figure S5K). Although these results are purely predictive, they may explain increased NETosis of FLNA^Q^ neutrophils (Figure 7K-L). ICAM1+ neutrophils have been shown to be more prone to ICAM1-RhoA-mediated NETosis, which relies on Rho kinase dependent actin remodeling (61). Along with this, we have found previously that FLNA^Q^ cells show mislocalization of p190Rho-GAP and increased levels of activated RhoA-GTP (9). In macrophages (MonoMac), enhanced Sema4d-Cd72 signaling may promote their phagocytic capacity (62), which could affect cytokine production and the removal of bacteria.

We lastly asked how FLNA editing recovers after acute colitis and compared control, acute DSS colitis (day ten) and samples from recovered animals (day 21). We found FLNA editing levels to be significantly reduced during acute disease (Figure 7O). Notably, humans showed the same dynamic with reduced FLNA editing levels in intestinal biopsies from patients suffering from acute UC, as compared to healthy controls or patients in remission (Figure 7P). Further, editing levels of mice and humans were most pronounced in the distal colon (Figure 7O-7P), highlighting the strong conservation of FLNA editing functions across species.

Together, we found that mice expressing only edited FLNA^R^ were highly protected from colitis, while a lack of FLNA editing rendered them more sensitive. DSS resistance is transplantable and reproducible in conditional, myeloid cell-specific FLNA^QiR^ LysMcre^+/−^ animals. Targeted introduction of an edited FLNA^R^ state in myeloid cells reduces intestinal inflammation and colitis severity, likely via its influence on neutrophils and inflammatory properties of macrophages. Our data also show a similar dynamic of FLNA editing during acute colitis and recovery in mice and human and thus highlight the therapeutic potential of targeted FLNA^R^ induction in IBD.

## Discussion

We show that the FLNA editing state is a key determinant of colitis severity. Physiologically, FLNA is strongly edited in the distal colon, precisely in fibroblasts, smooth muscle- and endothelial cells, and this is lost during acute colitis in mice and humans. Accordingly, mutant mice expressing only edited FLNA^R^ were DSS-resistant, whereas unedited FLNA^Q^ animals were hypersensitive. A fixed FLNA editing state widely affected transcriptional states of intestinal structural- and immune cells and altered the microbiome composition. Using cohousing-, depletion-, transplantation- and transgenic mouse approaches we find that protection via FLNA^R^ is largely caused by its immunomodulatory properties in myeloid cells, likely neutrophils and macrophages. Our study thus reveals novel insights into the modulating properties FLNA editing states in the colon and highlights the potential benefit of targeted, site-directed FLNA A to I editing (SDRE) in myeloid cells as a novel therapeutic approach for IBD.

Mice with a fixed FLNA^Q^ or FLNA^R^ state exhibit profound differences in colon epithelial- and CD8+ T cell transcriptomes. Together with a potentially protective, immunomodulatory microbiome with more butyrate (34) in FLNA^R^ mice, it was tempting to speculate that FLNA^R^ affected colitis susceptibility via the crosstalk between epithelial- and T-cells with the microbiota. This was further supported by a differential expression of defensins (*Reg3b* and *Reg3g*) between FLNA^R^ and FLNA^Q^ ECs and CD8+ T cells. *Reg3b* and *Reg3g* are protective in different settings of intestinal inflammation (19, 63) and typically get induced by Toll-like receptor activation (64). However, in two different experimental approaches, we could not prove a causal connection of microbiome composition and DSS-sensitivity, suggesting that distinct microbiomes are likely a consequence of genotype-specific host intrinsic changes. In spite of FLNAs importance for endothelial permeability (65) and observed differences in claudin and Mucin-4 expression, editing states did not impact epithelial barrier integrity. As the contribution of different claudins and Mucin-4 to IBD pathogenesis are still poorly understood (21), we conclude that FLNA editing modulates specific tight junction components, but cumulatively does not alter barrier leakiness.

Aside from its essential functions as cytoskeleton cross-linker, FLNA interacts with master regulators of cellular signaling like Syk or R-Ras (1). Genetic fixation of an unedited FLNA^Q^ or edited FLNA^R^ state resulted in the differential expression of genes involved in proliferation-, differentiation and survival signaling like AP-1 complex components (*Jun*, *Fos* and *Atf3*) (66) or *Hprt* (30) across different cell types, indicating that FLNA editing may adapt cellular responses to growth factors or cytokines. Importantly, these effects were prominent in structural cells, but also present in neutrophils during colitis, indicating cell-type independent features of edited FLNA. Syk promotes STAT1 activation (67) and as unedited FLNA^Q^ neutrophils upregulated genes associated with inflammation like *Ly6e, Tnfaip2* and *Stat1* itself (43), FLNA editing might suppress STAT1 activation. FLNA dependent Syk-regulation downstream of the TCR (68) may also explain transcriptional differences in the CD8+ T cell compartment. However, systemic CD8+ T cell depletion did not abrogate protection in FLNA^R^ animals and protection was driven by FLNA^R^ in Lysozyme-expressing, i.e. myeloid cells.

When investigating physiological FLNA editing, we found high levels only in the distal colon and could attribute this largely to endothelial cells and a cell fraction containing fibroblasts and SMCs. Of note, fibroblast/SMC and endothelial cell isolations were done from the whole colon and not only from the distal part, which may account for discrepancies with regards to exact editing frequencies between whole distal colon tissue and sorted cells. A limitation of our study is the lack of an endothelial cell population in our single cell sequencing dataset, caused by differential technical requirements to obtain healthy epithelial, endothelial and immune cell fractions. However, we can exclude a strong contribution of endothelial FLNA^R^ to the DSS resistance phenotype, due to its transplantability and its prevalence in FLNA^QiR^ LysMCre^+/−^ mice, which exclusively express edited FLNA^R^ in myeloid cells. Another limitation of the study is a lack of direct comparisons of FLNA^R^, FLNA^Q^ and, variably edited, wildtype mice. This was not feasible due to the huge impact of breeding and cage effects on intestinal homeostasis and - even more pronounced - DSS susceptibility. In our experience and according to the literature, littermate breeding is a critical requirement in studies using murine IBD models, which are particularly sensitive to cage-, microbiota- and genetic effects (69–71). While we would have preferred a setup that allows to draw conclusions about fixed editing states compared to physiologic editing, it is not possible to obtain littermates from all three genotypes. We thus refrained from including WT animals in most of our setups and compared the two fixed states of FLNA, which - we believe - is more accurate.

Since immune cells - in colon, blood and bone marrow - only exhibit low-grade editing in all tested conditions, our finding that protection from colitis is driven by myeloid FLNA^R^ appeared counterintuitive at first. However, FLNA impacts cell migration and adhesion by regulating surface integrin expression and by linking integrins to the actin cytoskeleton (72). While FLNA deficiency is compensated by FLNB in many cell types (73), FLNA deficiency rendered specifically neutrophils more adhesive, by enhancing their interaction with integrins (74). While we found reduced numbers of infiltrating neutrophils in the intestinal tissues of protected FLNA^R^ animals, we did not find a defect in FLNA^R^ neutrophil migration *in vitro*. In contrast, we found that FLNA^R^ neutrophils were slightly faster during at the onset of chemotaxis assays (1h). However, in later phases, this subtle defect was masked by the extreme temporal dynamics neutrophils develop at high population-densities (so-called swarming behavior) (75). Hence, it seems unlikely that such subtle differences impact pathology.

In neutrophils, FLNA was also shown to regulate reactive oxygen production and neutrophil extracellular trap formation (74), key antibacterial effector functions and drivers of tissue damage (76). We found protective FLNA^R^ neutrophils produced less and FLNA^Q^ neutrophils more NETs *in vitro*. Along with this, FLNA^R^ neutrophils showed downregulation of *Lmnb1*, which has been shown to critically influence nuclear envelope rupture during NET formation (77, 78). Counterintuitively, FLNA^R^ neutrophils appeared slightly more reactive to TLR stimuli. However, even though we did not find relevant transcriptional changes in macrophages, FLNA^R^ BMDMs showed an impaired capacity to produce IL-6, IL-1β and TNF upon TLR stimulation *in vitro*. Notably, macrophages, more than neutrophils, shape the inflammatory microenvironment of the colon (79, 80), which in turn is strongly influencing neutrophil NETosis (56, 57). Furthermore, macrophage migration in dense environments was shown to rely on FLNA and targeted reduction of FLNA impairs macrophage functions in an atherosclerosis model (81, 82). We thus propose a model in which, upon targeted editing in myeloid cells, FLNA^R^ in macrophages dampens their inflammatory responses to the microbiota upon DSS treatment, which in turn leads to a reduced recruitment of FLNA^R^ neutrophils. The already reduced inflammatory milieu and the intrinsic feature of FLNA^R^ neutrophils to be less prone to NETosis then mediates tissue protection and resistance to DSS induced colitis. This protection may be further amplified by a protective microbiome and anti-inflammatory transcriptional state of the epithelium in a background with a complete FLNA^R^ state. Reduced editing is found in humans during acute colitis, which increases upon remission. Thus, due to the high conservation of FLNA editing in terms of local levels and changes thereof during acute disease in mice and humans, SDRE of FLNA in myeloid cells may be exploited as a future therapeutic intervention for immunomodulation in settings of acute intestinal inflammation.

## Materials and methods

### Human material

Human intestinal biopsies were collected after written consent was obtained during regular colonoscopies at the Division of Gastroenterology and Hepatology of the Medical University of Vienna. In total biopsies of 21 patients were collected: samples of nine healthy donors, six active colitis patients (UC) and six colitis patients in remission, each with a proximal and distal biopsy. Clinical data and treatment regimens are summarized in Table S7. This study was conducted under the Health Research Authority of the Ethics Committee of the Medical University of Vienna, ethics approval number (“Ribonucleic acid variation in the intestine”, EK-Nr: 1692/2020, voted positively on 17. Sept. 2020, granted to CG).

### Mice

All animal experiments in this study were performed in compliance with the guidelines of the Ethical review committee of the Medical University of Vienna and national laws (approval numbers issued by the Austrian authorities: BMWFW-66.009/0260-WF/V/3b/2016 and 2021-0.203.240). All animals were used at an age of 8-12 weeks.

FLNA^Q^, FLNA^R^, IL10 knockout (KO), Vav-Cre and LysMCre mice were previously described (8, 9, 36, 51, 83). IL-10 KO mice were interbred with FLNA^R^ and FLNA^Q^ animals to obtain littermates of FLNA^WT^-IL-10^+/+^, FLNA^WT^-IL10^−/−^, FLNA^Q^-IL10^−/−^ and FLNA^R^-IL10^−/−^.

Inducible FLNA^QiR^ mice were generated for this study by Taconic Biosciences (former Cyagen, New York, USA). This mutant strain harbors an unedited FLNA^Q^ locus and an adjacent sequence encoding for a pre-edited FLNA^iR^ sequence to be exchanged with the FLNA^Q^ locus (Figure 6H). More specifically, exons 30-47 (NCBI Reference Sequence: XM_011247549.3; ATG start codon in exon 2, stop codon in exon 47, based on Transcript: ENSMUST00000114299) were replaced with a cassette containing the *Flna* CDS encoding FLNA^Q^, flanked by loxP sites plus the mutant CDS (loxP-endogenous SA of intron 29-CDS of exon 30∼47-3’UTR-3*SV40 pA-FRT-Neo cassette-FRT-loxP-endogenous SA of intron 29-mutant CDS of Exon 30∼47). Thus, in these mice Cre-recombination replaces the FLNA^Q^ (last codon in exon 42 = CAG) with the FLNA^iR^ CDS (last codon in exon 42 = CGG). A targeted ES cell clone was injected into C57BL/6 albino embryos, which were re-implanted into CD-1 pseudo-pregnant females. Founder FLNA^QiR^ animals were identified by their coat color, and their germline transmission was confirmed by breeding with C57BL/6 females and subsequent genotyping of the offspring. FLNA^QiR^ Vav-Cre^+/−^ and FLNA^QiR^ LysMCre^+/−^ mice, with expression of pre-edited FLNA^iR^ in hematopoietic/endothelial cells or myeloid cells and FLNA^QiR^ littermate controls were generated by crossing FLNA^QiR^ and Vav1Cre^+/−^ or LysMCre^+/−^ animals, respectively. Of note, FLNA^QiR^ Vav1Cre^+/−^ and and FLNA^QiR^ LysMCre^+/−^ mice were born at normal mendelian ratios, appeared healthy and did not show any obvious clinical phenotypes. All mice were bred on a C57BL/6 background and maintained in the specific pathogen-free animal facility of the Medical University of Vienna (22°C,12h light/12h dark cycle), with free access to food and sterile water.

### Murine colitis models

DSS-colitis was induced in 8-10-week-old mice by adding 2-2.5% DSS in the drinking water for seven days, followed by three days with regular drinking water. Body weight and health status of the animals were monitored throughout the experiment. Mice were euthanized on day five or ten by intraperitoneal Ketazol/Rompun overdose. At the endpoint, colons were isolated, measured and prepared for histology, RNA isolation, FACS analysis or sorting.

The development of spontaneous colitis in FLNA^WT^-IL-10^+/+^, FLNA^WT^-IL10^−/−^, FLNA^Q^-IL10^−/−^ mice was assessed by monitoring the bodyweight twice per week after weaning. All animals were euthanized at the time of onset of colitis symptoms at nine-ten weeks of age. Bodyweight and colon length were measured after sacrifice and the histological sections of colon rolls were scored for colitis parameters.

### Microbiome depletion

For antibiotic treatment, mice were randomly assigned to two groups. One group received a cocktail of antibiotics (1mg/mL ampicillin, 1mg/mL vancomycin, 1mg/mL streptomycin and 1mg/mL neomycin) in drinking water for a total of 21 days, while the control group received drinking water only. During antibiotic treatment, water bottles from both groups were replaced every 2-3 days. Fresh fecal pellets were collected from every mouse (antibiotic treated (Abx) and non-antibiotic treated) at the end of the antibiotic treatment and immediately placed in a sterile tube. Half of the fecal sample was transferred into a tared Eppendorf tube, weighed, and homogenized in sterile phosphate buffered saline (1xPBS) containing 20% glycerol for final concentration of 10mg/mL with the help of Eppendorf pestle, and afterwards stored at −80°C. The remaining half was immediately frozen in dry ice and stored at −80°C for nucleic acid extraction and 16S rRNA amplicon sequencing analyses.

### Bone marrow transplantation

Mice got lethally irradiated (2 x 6Gy) at an age of seven weeks with a YXLON Maxishot (YXLON International GmbH), with a 3hrs break between each dose. In parallel, bone marrow cells were isolated from femurs and tibias from 8-week old donors. Epiphyses were cut, and bone marrow was flushed out with RPMI-1640 (Gibco) medium using a 1ml syringe (with 27 G needles), filtered through a 70µm strainer, counted and resuspended in 100µl 1xPBS for retro-orbital injection into recipient mice. 5 × 10^6^ bone marrow cells were injected, four hours after the final irradiation with mice maintained under light isoflurane anesthesia (2% isoflurane 2L/min O_2_). Mice were housed for 8 weeks to allow for complete reconstitution before the start of the respective experiments.

### Colitis score – histological preparation and scoring

Pathological scoring was performed on day ten post colitis induction by DSS. Colons (without cecum) were dissected, measured (length), cut open longitudinally and rolled up starting from the distal colon. Colon rolls were fixed in 4% paraformaldehyde in 1xPBS for a minimum of 24hrs. The samples were further paraffin embedded and sectioned at 5μm thickness. Sections were stained with hematoxylin-eosin and scored for mild, moderate or severe colitis according to the following parameters: Crypt damage (0 = none, 1 = 1/3 loss of basal, 2 = 2/3 loss of basal, 3 = complete destruction), inflammatory cell infiltration (0 = none, 1 = mucosal, 2 = mucosal and submucosal/moderate, 3 = mucosal and submucosal/severe), epithelial erosion and ulceration (0 = none, 1 = light submucosal erosion, 2 = partial submucosal erosion, 3 = confluent submucosal erosion) and thickening (0 = none, 1 = mild/focal, 2 = partial, 3 = complete). Each score was multiplied by 0 (= 0%), 1 (= 1-25%), 2 (= 26-50%) or 3 (= 51-100%) depending on the % of the area involved. Scoring was performed blinded by a trained lab technician. Proximal, middle and distal colon was first scored separately and then a combined total score was calculated.

### Colon single cell preparations for single cell sequencing, flow cytometry and FACS sorting

The large intestine was removed from freshly sacrificed animals, cut open longitudinally and fecal contents were removed by vigorous shaking in PBS. For single cell sequencing and FACS sorting of immune cells, colons were dissected as previously described (53). Briefly, they were incubated at 37°C for 15min in 25mL IEL solution (2% FCS, 10mM HEPES, 1% L-glutamine, 1% Pen/Strep, 1mM DTT, 1mM EDTA in PBS) at 250rpm. 1/10 of the cells dissociated using EDTA were directly lysed in RNA Lysisbuffer for the sequencing of epithelial cells. Tissues were then briefly rinsed with PBS, transferred to 50mL falcons containing ceramic beads and 25mL collagenase solution (2% FCS, 10mM HEPES, 1% L-glutamine, 1% Pen/Strep, 1 U/mL DNAse I, 1mg/mL collagenase A in RPMI (Gibco)) and incubated for 30min at 37°C rotating at 250rpm. Released lamina propria contents were then filtered through a 100μM cell strainer, washed with 1xPBS containing 2% FCS to remove remaining collagenase. Colon endothelial cells of wild type mice were isolated using another published protocol (84), with some modifications: Before colon dissection, anesthetized mice were perfused with 4mL of 1000U/mL Heparin at a perfusion rate of 2mL/minute. The endothelial cell enrichment step using CD31 MicroBeads was skipped and the obtained cells were directly stained as described below. For flow cytometry, colon single cells were incubated at 37°C for 20min in 4mL solution 1 (3% FCS, 20mM HEPES, 5mM EDTA, and 1mM DTT in RPMI (Gibco)) at 180rpm in a 6-well plate. Tissues were rinsed with 1xPBS, transferred to a fresh 6-well plate containing 4mL of digest1 solution (0.8mg/ml Dispase, 20mM HEPES, 2mM EDTA in RPMI) and again incubated at 37°C at 180rpm for 5min. Next, tissues were agitated vigorously in digest1 solution before transferring them into digest2 solution (0.1mg/ml Liberase TL, 0.5mg/ml DNAseI, 20 mM HEPES in RPMI). After another incubation for 20min, tissue and solution were transferred into a gentleMACS™ M tube and dissociated in an octo-gentleMACS dissociator (Myltenyi Biotech, Germany). Suspensions were then filtered through a 100μM cell strainer, washed with 1xPBS containing 2% FCS to remove remaining enzymes.

### Flow cytometry and cell sorting

For flow cytometry, cells were resuspended in antibody mixtures (all from BioLegend, fixable viability dye eF780, CD16/CD32 Fc block, CD45 PE TexasRed/AF700/PerCP Cy5.5, CD3 FITC/PE TexasRed, CD19 FITC/BV605, F4/80 FITC/PB, CD11b AF700/PECy7, Ly-6G PECy-7/FITC, CD8 BV510, CD4 PerCP Cy5.5, NK1.1 APC, Ly-6C BV510, CD11c PE TexasRed, BST2 PE, B220 AF700, CD103 APC, all from BioLegend) and stained for flow cytometry as previously described (85). For single cell sequencing, cells were resuspended in antibody mixtures, stained for 30min in the dark and viable CD45+ and CD45-cells were sorted using a FACS ARIA into 1xPBS/0.2% BSA. Sorted cells were then pooled, counted and fixed for 18hrs at 4°C in 1mL of freshly prepared fixation buffer (Fixation of Cells & Nuclei for Chromium Fixed RNA Profiling). On the next day, cells were pelleted, resuspended in freezing solution and stored at −80°C until processing for fixed single cell sequencing. Library preparation was performed using the Chromium mouse transcriptome probe set v1.0.1 protocol. Libraries were sequenced on an Illumina HiSeq 4000 and count matrices were generated using default parameters of the CellRanger version 7.1.0 software for the fixed assay. For the sequencing of immune cells, B cells (CD45+, CD19+), T cells (CD45+, CD19-, CD3+), neutrophils (CD45+, CD3-, CD19-, Ly-6G+, CD11b+) and a myeloid cell-enriched fraction (CD45+, CD3-, CD19-, Ly-6G-, CD11b+) were sorted directly into RNA lysis buffer. Endothelial cells and a fibroblast/SMC-enriched fraction, cells were stained with anti-CD31, anti-EpCAM and anti-CD45 antibodies and sorted into RNA lysis buffer (endothelial cells: CD45-, EpCam-, CD31+, fibroblast/SMC fraction: CD45-, EpCam-, CD31-).

### scRNA sequencing analysis

Quality control was performed per sample on the pre-filtered CellRanger output matrices. Data were imported as Seurat (v.4.3.0) or Scanpy AnnData (v.1.9.1) object and processed with custom R (v.4.1.0) or Python (v.3.8.12) scripts respectively throughout the analysis.

Cells with mitochondrial reads > 5 %, hemoglobin reads > 10%, or total UMI count < 1000 were filtered out. Only genes with at least three reads in three cells were retained. Doublet detection was performed with the SOLO adaption of SCVI and doubletFinder using default settings. Cell barcodes identified by either method were labeled as doublet. In addition, we used the SCVI latent space to identify clusters that are significantly enriched with doublet cells. All cells and clusters identified as doublet were excluded. In a last step we removed cells with gene counts in the lower 1% percent quantile. For integration the samples were first merged before configuring a SCVI model with the sample name as batch key and the genotype and treatment as categorical covariate. The model was trained with two hidden layers for the encoder, a negative binomial distribution for the gene likelihood (86) and otherwise default settings. A neighborhood graph (n_neighbors=50) was constructed on the latent space and used for Leiden clustering and uniform manifold approximation and Projection (UMAP) embedding with Scanpy. Cell type annotation of the Leiden cluster was performed using marker gene expression and supported by SingleR annotation with the ImmGen reference. Clusters with ambiguous marker gene expressions were subclustered using resolution parameters ranging from 0.1 to 0.2. For the final annotation, clusters with the same cell type label were merged and evaluated by computing marker gene expression with the Seurat implementation of the Wilcoxon test. The cell type annotation was transferred to additional WT samples using SCANVI (87). To measure the genotype effect on baseline or DSS treated cells we performed differential gene expression analysis with MAST. The analysis was restricted to cell types with at least 10 cells per group. The DEA results were ranked by the log2FC signed -log10 adjusted p-value for gene set enrichment analysis (32) with the MSigDB HALLMARK data base. Cell-cell interaction was inferred with CellChat v2 (59, 88).

### Distal colon bulk-Sequencing transcriptomics

Mice were treated with DSS or plain drinking water for five or seven plus three days, sacrificed and colons dissected and flushed with 1xPBS. The distal 1/3 of the colon was processed for RNA extraction. Tissue was homogenized using Circonia Beads (1mm) and a FastPrep®-24 homogenizer (VWR, Avantor). Total RNA was subjected to poly-A RNA enrichment (NEBNext® Poly(A) mRNA Isolation Module, NEB,) followed by stranded library preparation (NEBNext® Ultra™ Directional RNA Library Prep Kit for Illumina, NEB), multiplexed and sequenced on a NextSeq Illumina platform to a minimum of 30 Mio reads per sample. The 5-day time-point used 75bp single-end sequencing and the ten-day timepoint used 100bp of paired-end sequencing mode. Quality control and filtering of the raw reads was performed with FASTP. The reads were then mapped to the GRCm38 reference genome using the GENCODE M25 annotation with STAR (v.2.7.9a). The BAM files were converted to a unified count matrix with featureCounts. For differential expression analysis (DEA) the data were first subset for each comparison. Lowly expressed genes were then filtered out by only retaining genes with at least five counts in two samples in either group. The count matrix was then processed with the Limma “voomWithQualityWeights” function before applying the standard Limma workflow for DEA. The DEA results were ranked by the log2FC signed -log10 adjusted p-value for gene set enrichment analysis (32) with the MSigDB HALLMARK data base.

### Determination of FLNA Q/R site editing levels

For the determination of FLNA editing ratios in whole tissue samples, total RNA was isolated from homogenized human biopsies or mouse tissue with TRIfast reagent using the manufacturer’s protocol. After DNase I treatment, cDNAs were synthesized using LunaScript Reverse Transcriptase and random hexamer primers. A FLNA cDNA fragment spanning spliced exons 42–43 was amplified by PCR, gel-eluted, and Sanger sequenced to check editing levels (human and mouse primers used for amplification and sequencing see Supplementary material table). For human biopsies, two samples per location (2xproximal and 2xdistal colon) were obtained and analyzed. In cases where the Sanger sequencing chromatogram was of inferior quality (background of all other bases > 5-10%) in one of the samples, only the high-quality sample was used for statistical analysis. In cases where both proximal or both distal samples had a good and comparable quality of their Sanger sequencing chromatograms, their average was used to calculate editing frequencies.

For the detection of FLNA editing levels in sorted cell populations RNA was extracted using the NucleoSpin 96 RNA Core Kit (Macherey-Nagel). CarrierRNA (Macherey-Nagel) was added to samples with cell counts <10^4^ cells. cDNA was synthesized using the LunaScript RT Mastermix (NEB) and then subjected to a 2-step PCR: In a first round, Flna editing site primers (amplification of the region around the editing site) and Q5 High-Fidelity DNA Polymerase (NEB) were used to perform 25 cycles. 2µl of this PCR served as a template for PCR2: Using the ampSeq barcoding primers for 15 cycles, thereby adding Illumina-compatible adapters with dual-indices. PCR products were gel-eluted using the Monarch DNA Gel Extraction Kit (NEB) followed by quantification by Qubit Fluorometry (Thermo Fisher Scientific). A maximum of 96 samples were pooled into one multiplex for sequencing on an Illumina Nova- or Next-Seq instrument. After quality control using fastqc and multiqc, trimming was performed in two consecutive steps with cutadapt (--nextseq-trim=30 --minimum-length=10 (a) nextseq-trim; then: -a -A --minimum-length=10), resulting in, on average, 58.000 processed reads/sample (150bp paired-end). Reads were mapped using Hisat2 and editing ratios of each sample were calculated using Jacusa2.

### RT-qPCR

RNA was isolated with NucleoSpin RNA Core Kit including lyophilised carrierRNA for sorted cells (Macherey-Nagel). For real-time PCR assays, cDNA synthesis was performed using the LunaScript cDNA Synthesis Kit (NEB), according to manufacturer’s protocol. Real-time PCR was performed with SYBR Green Master Mix reagents (Avantor, VWR) on a StepOnePlus Real-Time PCR System (Applied Biosystems). Mouse primers used for amplification of cytokines, *Flna*, *Adar* and *Adarb1* are found in the supplementary material table. Transcript levels were normalized to *Gapdh*.

### Legendplex

Tissue cytokines and chemokines were measured using the LEGENDplex^TM^ Mouse Anti-Virus Response Panel (13-plex) with V-Bottom Plate (BioLegend). Samples were prepared according to the manufacturer’s protocols and analysed by flow cytometry. Data analysis was performed using the LEGENDplex data analysis software.

### Assessment of intestinal epithelial barrier permeability

To test the intestinal permeability *in vivo*, FLNA^Q^ and FLNA^R^ mice were starved for four hours followed by oral gavage with 0.6mg/g body weight FITC-dextran 4000. Serum was collected four hours post gavage and concentrations were measured by fluorometry (Excitation: 485 nm, emission: 528 nm, EnSight, Perkin Elmer).

Barrier permeability was additionally assessed *ex vivo*. Colons of sacrificed FLNA^R^ and FLNA^Q^ mice were flushed with 1xPBS and filled with 1ml FITC-dextran 4000 (1mg/ml in PBS). Colons were closed from both sides and incubated at room temperature for 40min. Following incubation, colons were fixed and prepared for paraffin sectioning. Proximal and distal colons were sectioned horizontally into two different segments each and FITC positive cells were visualized using fluorescence microscopy after DAPI staining in an Olympus VS120-S6 fluorescence microscope.

### Fecal pellet output measurement

FLNA^Q^ and FLNA^R^ mice were kept separated by genotype in cages (4 mice/cage) for 24hrs with paper towel sheets instead of regular bedding. After 24hrs sheets were removed and air dried for another 24 hours and the pellet number and dry weight was assessed.

### DNA isolation from fecal pellets and 16S rRNA gene amplification, sequencing and analysis

DNA was isolated with the QIAamp Fast DNA Stool Kit according to manufacturer’s instructions in an automated manner using a QiaCube Symphony at the Joint Microbiome Facility (Vienna, Austria). Amplification of bacterial and archaeal 16S rRNA genes from DNA samples was performed with a two-step barcoding approach. In the first-step PCR, the primers 515F and 806R including a 5’-head sequence for 2-step PCR barcoding, were used. PCRs, barcoding, library preparation and Illumina MiSeq sequencing were performed by the Joint Microbiome Facility (Vienna, Austria) under project number JMF-2005-3. First-step PCRs were performed in triplicates (12.5μl vol. per reaction) with the following conditions: 1XxDreamTaq Buffer (Thermo Fisher), 2mM MgCl_2_ (Thermo Fisher), 0.2mM dNTP mix (Thermo Fisher), 0.2µM of forward and reverse primer each, 0.08mg/ml Bovine Serum Albumin (Thermo Fisher), 0.02 U Dream Taq Polymerase (Thermo Fisher), and 0.5µl of DNA template. Conditions for thermal cycling were: 95°C for 3 min, followed by 30 cycles of 30 sec at 95°C, 30 sec at 52°C and 50 sec at 72°C, and finally 10 min at 72°C. Triplicates were combined for barcoding (8 cycles). Barcoded samples were purified and normalized over a SequalPrep™ Normalization Plate Kit using a Biomek® NXP Span-8 pipetting robot (Beckman Coulter), and pooled and concentrated on columns (Analytik Jena). Indexed sequencing libraries were prepared with the Illumina TruSeq Nano Kit and sequenced in paired-end mode (2× 300 bp; v3 chemistry) on an Illumina MiSeq following the manufacturer’s instructions. The workflow systematically included four negative controls (PCR blanks, i.e., PCR-grade water as template) for each 90 samples sequenced.

Amplicon pools were extracted from the raw sequencing data using the FASTQ workflow in BaseSpace (Illumina) with default parameters. Input data was filtered for PhiX contamination with BBDuk. Demultiplexing was performed with the python package “demultiplex” allowing one mismatch for barcodes and two mismatches for linkers and primers. Barcodes, linkers, and primers were trimmed off using BBDuk with 47 and 48 bases being left-trimmed for F.1/R.2 and F.2/R.1, respectively. DADA2 R package was used for demultiplexing amplicon sequencing variants (ASVs) using a previously described standard protocol. FASTQ reads were trimmed at 150nts with allowed expected errors of two. Taxonomy was assigned to 16S rRNA gene sequences based on SILVA taxonomy (release 138) using the DADA2 classifier. Amplicon sequence libraries were analyzed using the vegan (v2.4.3) and phyloseq (v1.30.0) packages of the software R. DESeq2 (v1.26.0) implemented in phyloseq was used to determine statistically significant differences in ASV abundances between groups of mice. Only ASVs that had ≥10 reads were considered for comparisons by DESeq2 analyses. All statistical analysis on microbiome data was carried out with the software R (R 4.0.2), and statistical tests and p-values are indicated in the figure legends. The 16S rRNA gene sequences were deposited in the NCBI Sequence Read Archive (BioProject ID: PRJNA1051489).

### Short chain fatty acid measurements

Fecal pellets (average weight 21.62 ± 4.90 mg) were suspended in 0.5 mL milli-Q water (Milli-Q, Merck Millipore, Burlington, MA, US) and homogenized with a pellet mixer (VWR, Radnor, PA, US). Homogeneous suspensions were briefly vortexed at maximum speed after adding 1 mL milli-Q water. The supernatant after centrifugation samples for 5min at 10,000*g* was filtered over a 0.2µm filter (0.2 µm sterile syringe filter, diameter 26mm, Minisart, Sartorius, Göttingen, Germany). Filtered samples were analyzed on a 930 Compact IC Flex instrument equipped with an 858 Professional Sample Processor with extended MiPuT (Micro Metrohm intelligent Partial Loop Injection Technique), a Metrohm CO_2_ Suppressor for inline bicarbonate removal, a Metrosep Organic acids 250/7.8 column, a Metrosep Organic acids Guard/4.6 guard column and a 850 IC conductivity detector (Metrohm, Herisau, Switzerland). Data was analyzed using the software R (R 4.0.2), and statistical tests and p-values are indicated in the figure legends.

### Quantification of fecal bacterial density by flow cytometry

Microbial loads from mouse fecal samples preserved in 1xPBS containing 20% glycerol were determined using flow cytometry and counting beads as detailed below. Samples were diluted 10 to 500 times in 1xPBS. To remove any additional debris from the fecal matrix, samples were transferred into a flow cytometry tube by passing the sample through a snap cap containing a 35μm pore size nylon mesh. Next, 500μL of the microbial cell suspension was stained with SYTO™ 9 (0.5μM in DMSO) for 15min in the dark. The flow cytometry analysis of the microbial cells present in the suspension was performed using a BD FACS Melody (BD Biosciences), equipped with a BD FACSChorus software (BD). Briefly, background noise of the machine and of PBS was detected using the parameters forward scatter (FSC) and side scatter (SSC). Microbial cells (no beads added) were then displayed using the same settings in a scatter plot using the forward scatter (FSC) and side scatter (SSC) and pre-gated. Singlets discrimination was performed. Absolute counting beads added to each sample were used to determine the number of cells per mL of fecal slurries by following the manufacturer’s instructions. Fluorescence events were monitored using the blue (488 nm – staining with SYTO™ 9 and CountBrightTM beads) and yellow-green (561 nm - CountBrightTM beads only) optical lasers. The gated fluorescence events were evaluated on the forward–sideways density plot, to exclude remaining background events and to obtain an accurate microbial cell count. Instrument and gating settings were identical for all samples (fixed staining–gating strategy).

### Bone marrow neutrophil isolation and bonemarrow derived macrophage differentiation

Bone marrow neutrophils were isolated from femurs of 8-10 week-old mice using the Neutrophil isolation kit, mouse (Miltenyi Biotec) following the provided protocol. In brief: femurs were flushed using a 25-gauge needle and BM cells were gently dissociated using a 1000µl pipette tip and filtered through a 30µm cell strainer. Cells were then sequentially incubated with the antibody cocktail and the microbeads provided in the kit followed by negative selection on a QuadroMACS magnetic MACS separator using LS columns (both Miltenyi Biotec). For the preparation of bone marrow derived macrophages, bone marrow was flushed from femurs with PBS and cells were differentiated for 6 days in RPMI 1640 supplemented with 10% FCS, 1% pen-strep and 10% L929 conditioned medium.

### Neutrophil DNA release assay

DNA release as a measure for neutrophil DNA trap release was performed as before (89). In short, 1×10^5^ BM neutrophils suspended in 100μl of Hank’s balanced salt solution with Ca^2+^ and Mg^2+^ (HBSS++) were seeded and then stimulated with 0.5μM of a23187, or with 0.1nM of phorbol 12-myristate 13-acetate (PMA) for 300 min at 37°C. For quantification of DNA release, 5μM of SYTOX Green was added and fluorescence was measured at 15 min intervals at excitation/emission of 485/520nm using the VarioSkan Lux plate reader (Thermo Fisher Scientific Inc).

### NETosis immunofluorescence

Colon sections (5μm) were analyzed for neutrophils (Ly6G) and NETs (CitH4 and DNA) by immunofluorescence staining as reported in (90). Sections were deparaffinized and rehydrated using xylene and decreasing concentrations of ethanol. After antigen retrieval in 0.5M citrate buffer, colon sections were blocked in 1xPBS with 5% bovine serum albumin and then stained for NETs with rat anti-mouse Ly6G for neutrophils (1:1000), rabbit polyclonal anti-CitH4 antibody (1:500), and secondary antibodies: Cy3-conjugated donkey anti-rat IgG and Alexa Fluor 647-labeled donkey anti-rabbit IgG (1:400), as well as Hoechst 33342 (1:1000) for the nuclear stain. Stained sections were imaged with an automated Axio Observer Z1 microscope (Carl Zeiss MicroImaging, Inc., Oberkochen, Germany) with a 20x objective and theTissueFAXS scan software. The tissue scoring of neutrophils and NETs was conducted manually by ordinal scoring based on 4 levels of signal frequency: 0–none, 1–few, 2–moderate, 3–high number of detectable cells.

### CD8+ T cell depletion

8-10 week old male mice received either anti-mouse CD8α antibody (CD8 YTS169; 500µg, *i.p.*) or the same volume of 1xPBS starting at 36h before DSS treatment start and then again on day 1 and 3 after DSS start (with 2% DSS in drinking water ad libido for 5 days). Mice underwent daily checkups including weight measurements.

### Neutrophil migration speed measurements

To distinguish cells by genotype in a microfluidic device, FLNA^Q^ and FLNA^R^ BM-Neutrophils were counted and stained with 10µM TAMRA and CFSE (Thermo Fisher Scientific) at 1000 cells/µl for 10mins at room temperature in the dark. Cells were washed three times and then mixed in equal numbers of FLNA^Q^ and FLNA^R^. To exclude dye-effects on the cells, dye-swap controls were included in every experimental set.

Microfabricated PDMS devices were manufactured as described in (91). The pillar forest had pillars of 3.8µm height, and 10µm width, in a square layout. The pore size between pillars was 5µm and the analysis area was 0.7mm x 2.5mm in size. PDMS devices were preincubated with RPMI with 2% BSA prior to use and loaded on one side with 50µM fMLP and on the other side with BMDNs (20.000 cell/µl). After 1 hour incubation, movies of stained cells in the pillar maze were acquired by time-lapse acquisition (90s intervals) using a Nikon Ti2E inverted widefield microscope with a plan Apo λ 20x/0.75 DIC 1 air PFS objective and a NIKON FX-format, monochrome CMOS sensor camera. During imaging, devices were kept in a custom-built climate chamber (37°C, 5% CO2, humidified). Spot analysis to calculate the mean cell speed was performed using the tracking tool of Imaris 9.9.1 software (Oxford Instruments). Only the first 40 frames (= 1 hour observation time) were used for cell tracking.

### Statistical analysis

Data were analyzed using GraphPad Prism 9.1 (GraphPad Software). All details about statistical testing of individual experiments are explained in the respective legends. In brief, data are represented as a mean +/− SD or SEM whenever indicated. Differences between two groups were tested using Student’s *t-*test (2-sided) or non-parametric Mann Whitney U test (for histological scores). For multiple group comparisons data were analyzed using one-way ANOVA or 2-way ANOVA (for weight-loss curves or editing time-courses) or Kruskal-Wallis test (for histological scores), post-hoc followed by a Tukey multiple comparison test. Statistical significance was attributed to p-values ≤ 0.05. Measurements were taken from distinct samples (unless specified: e.g. distal and proximal colon), except for mouse weight measurements, which were taken from the same individuals on different days.

## Materials

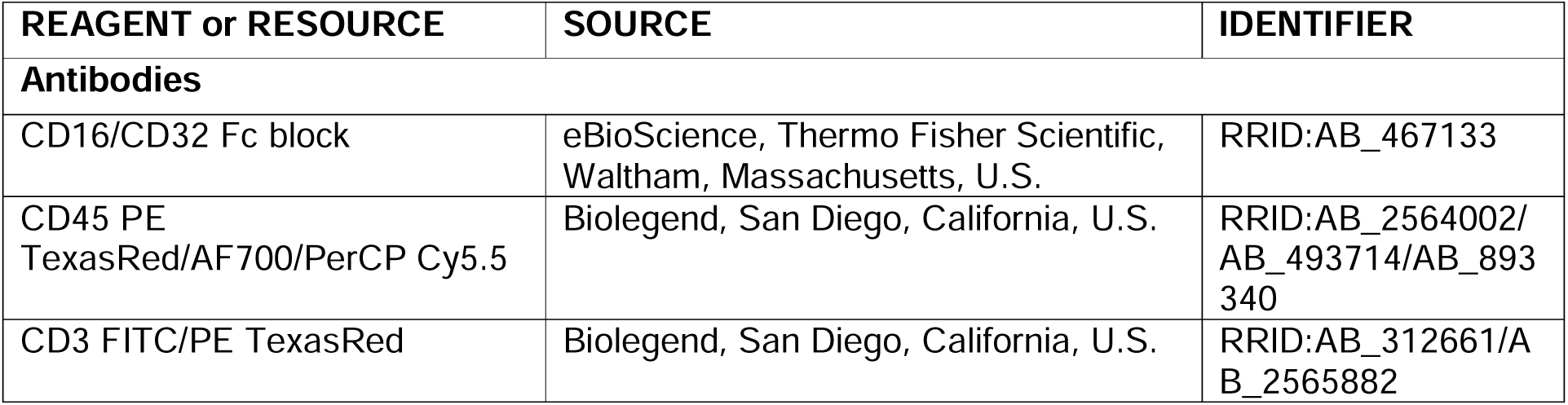
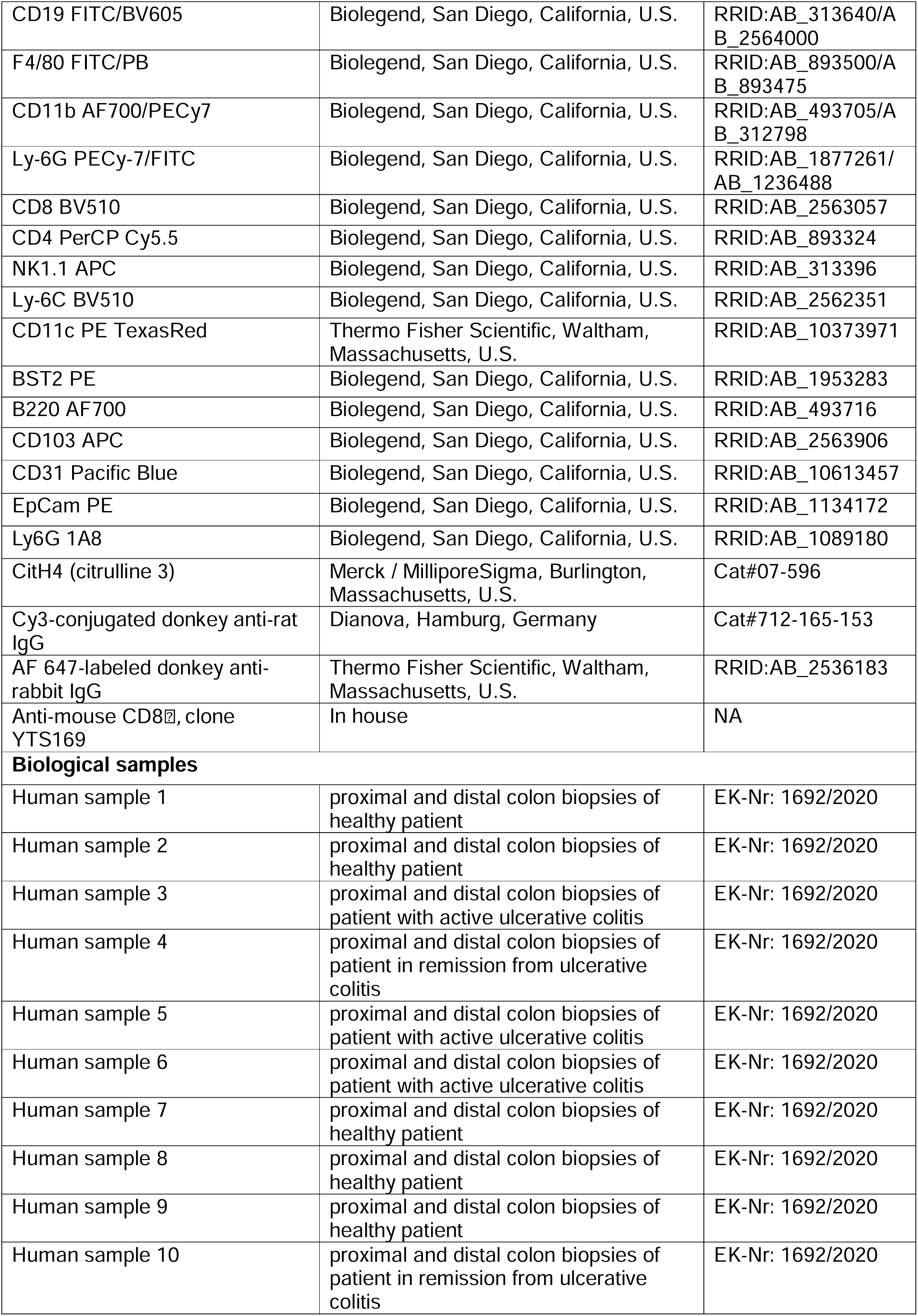
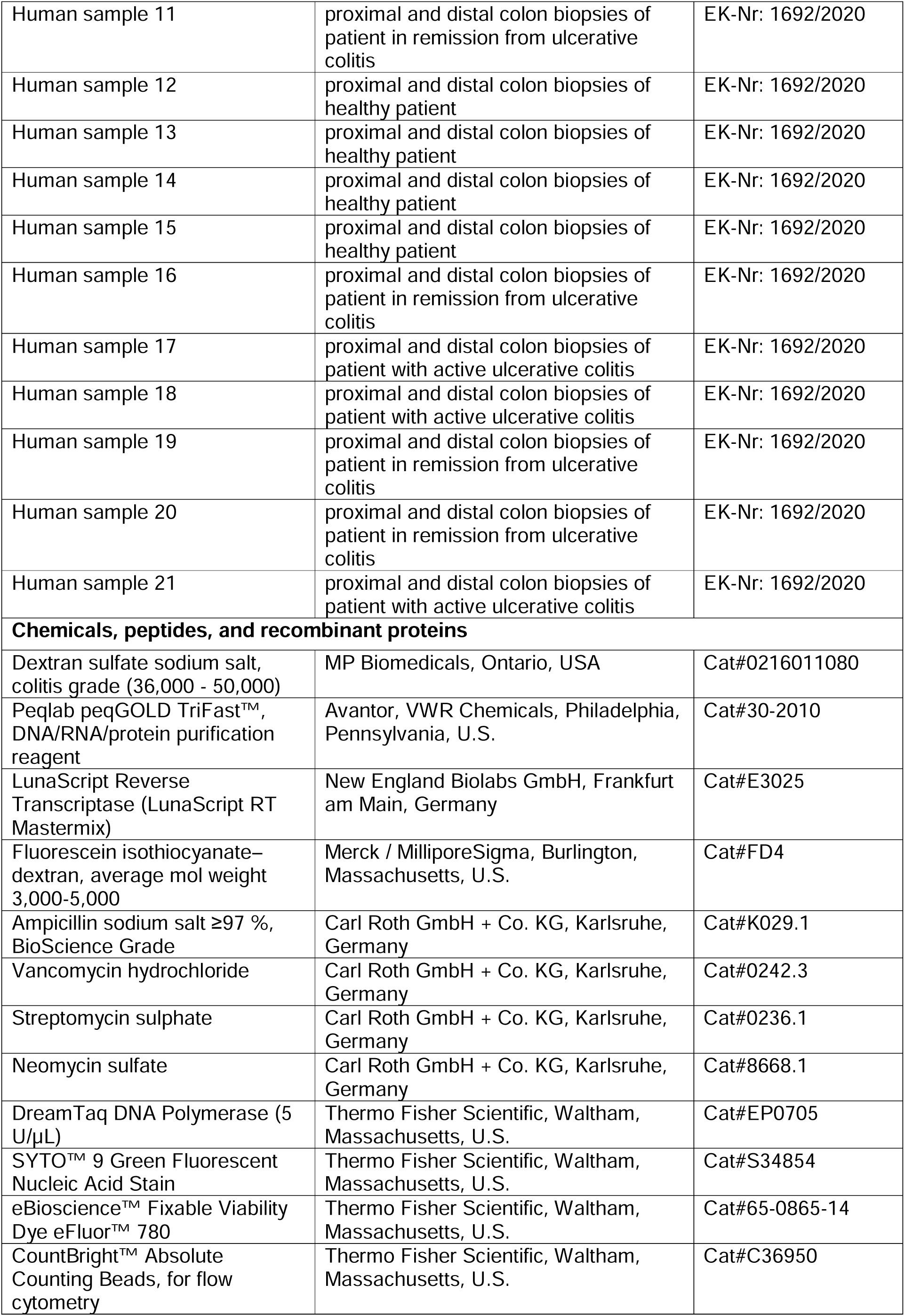
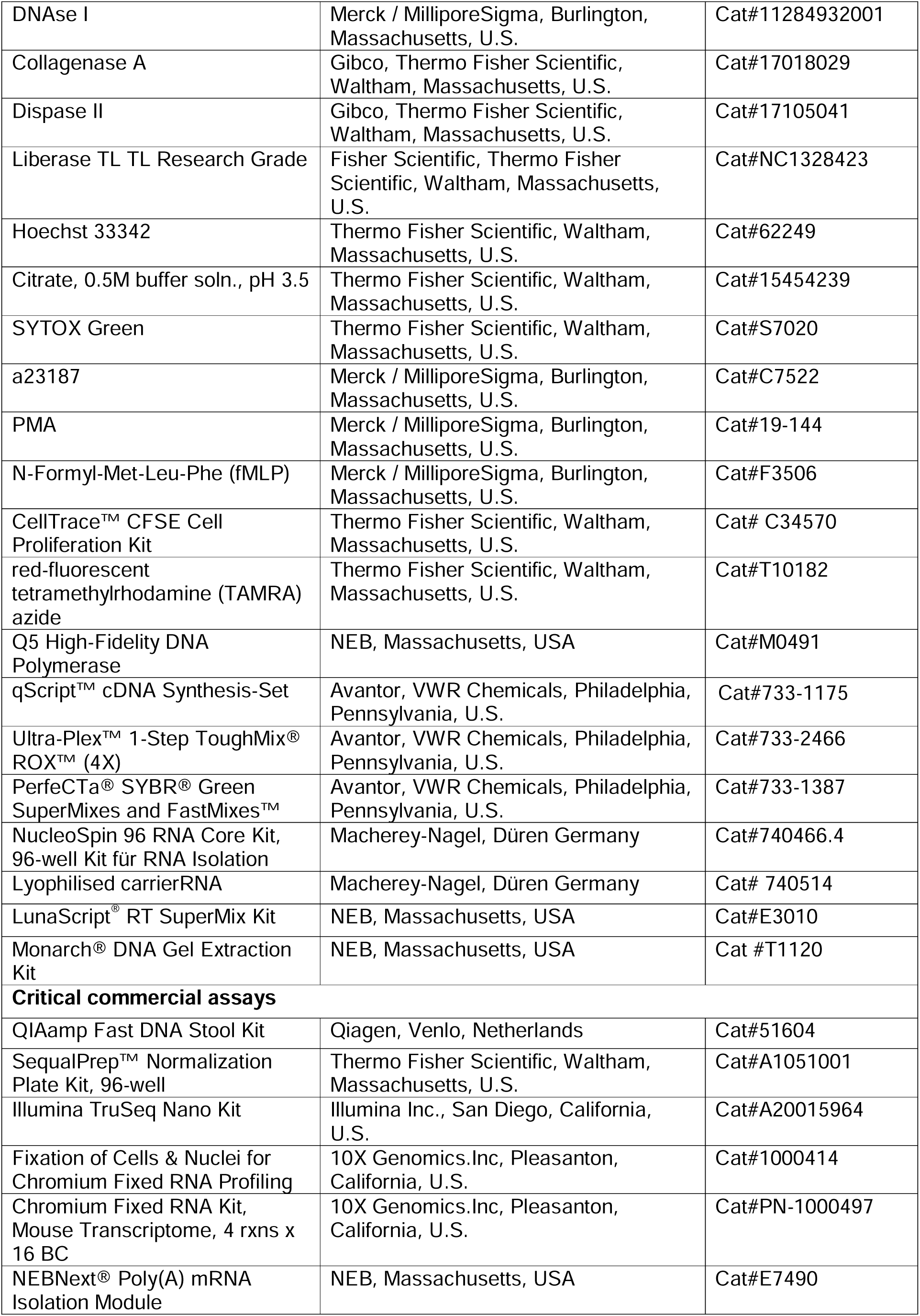
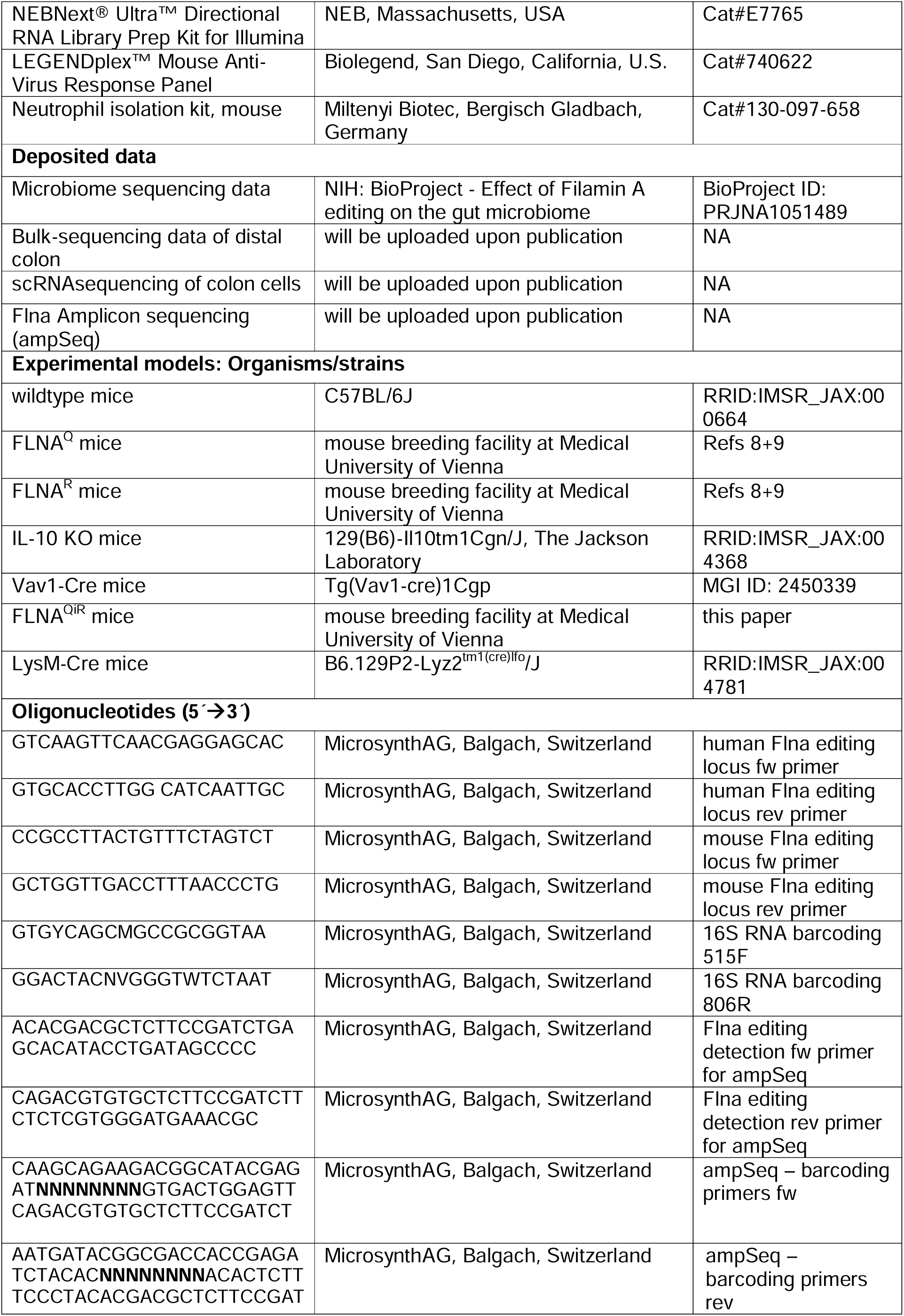
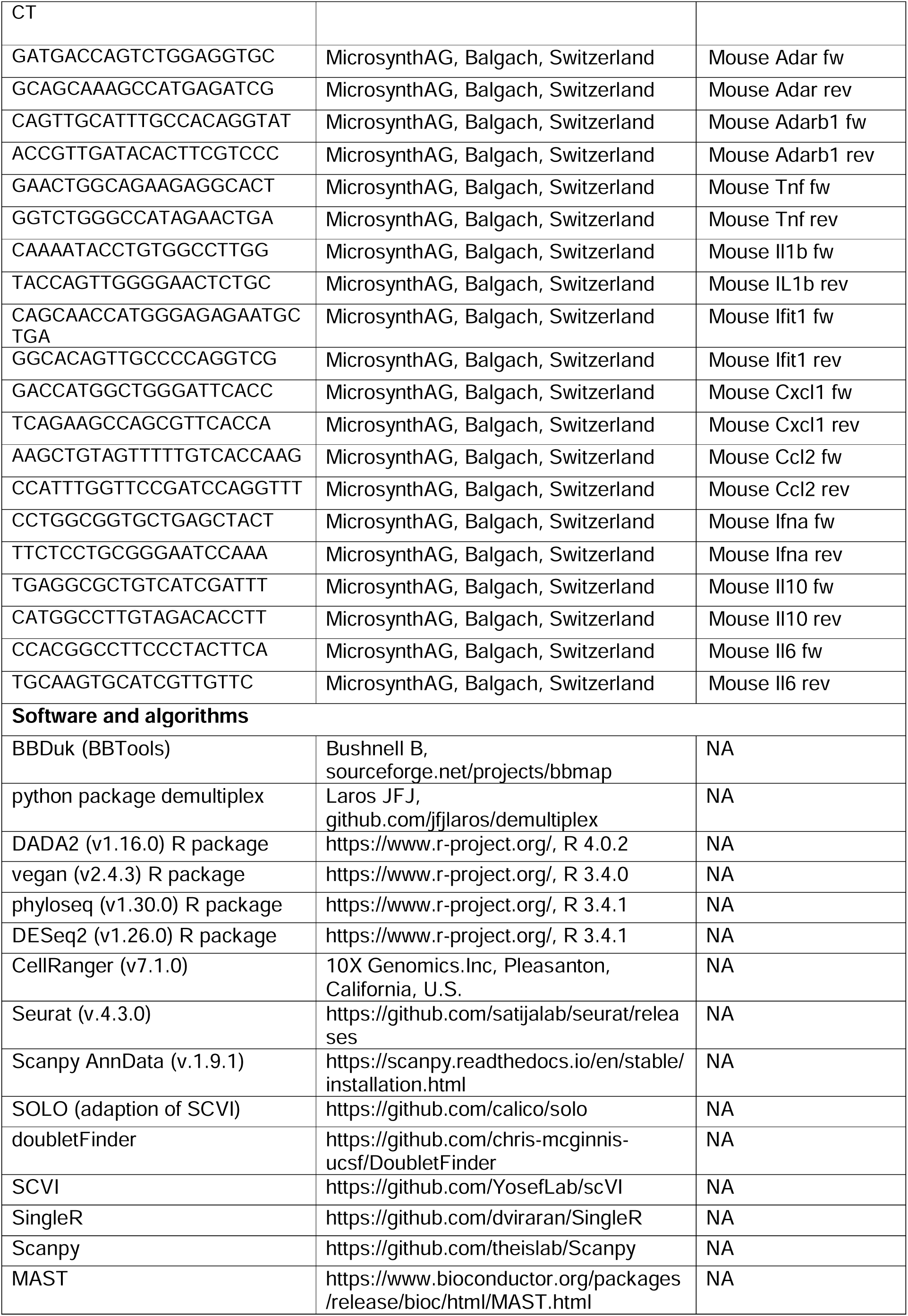
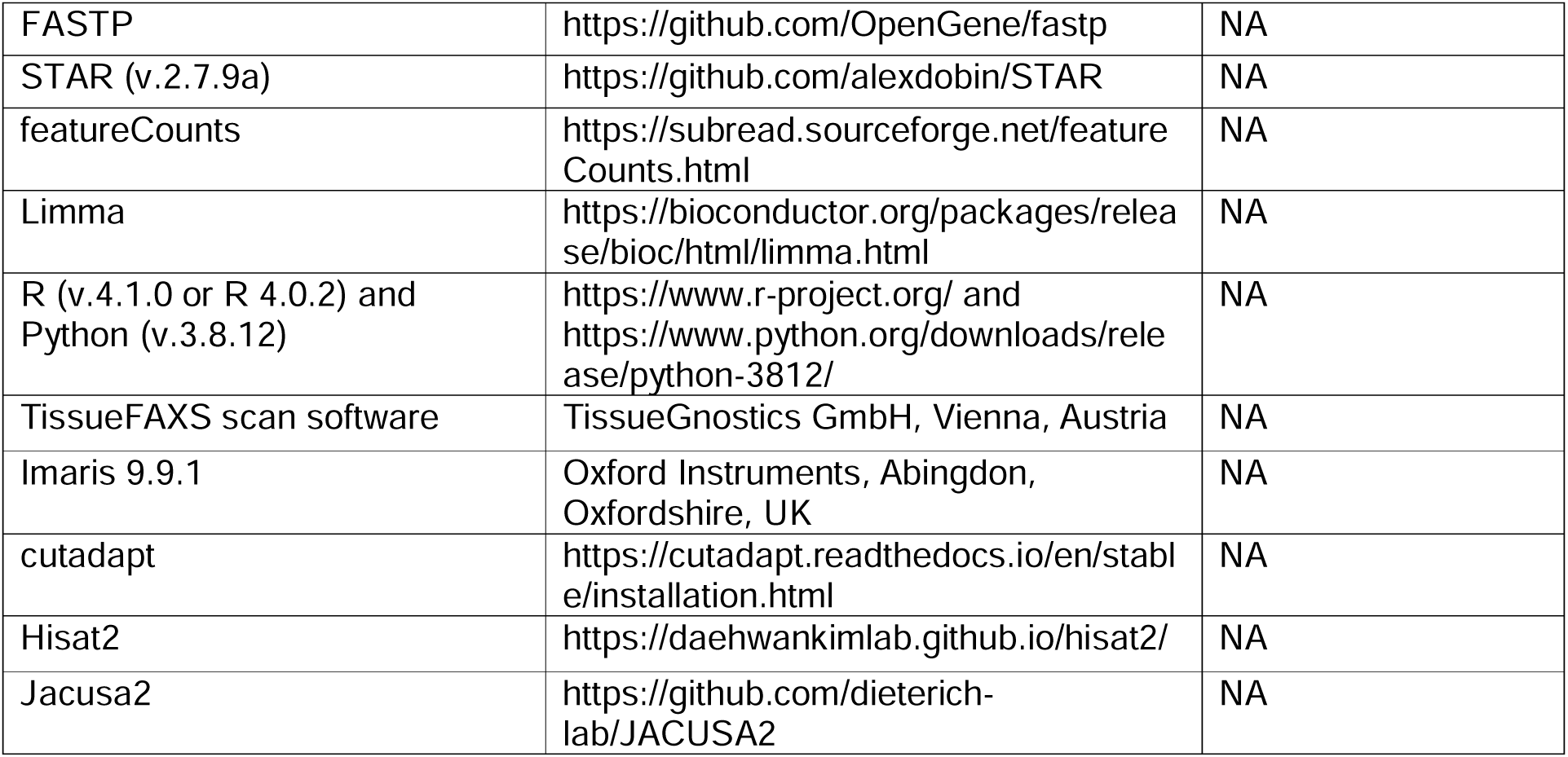

## Data availability statement

Transcriptomic data will be made available under GEO upon publication of this manuscript, all other data are available in the main manuscript or supplemental data. 16S rRNA gene sequences were deposited in the NCBI (https://www.ncbi.nlm.nih.gov/) Sequence Read Archive (SRA) under BioProject ID PRJNA1051489.

## Acknowledgements

RNA and single cell RNA sequencing was performed by the VBCF (Medical University Vienna Core facility) and the BSF (Biomedical Sequencing Facility at CEMM, Vienna) and cell sorting as well as flow cytometry was performed at the Core Facility Flow Cytometry and Imaging (Medical University of Vienna). We thank Jasmin Schwarz, Gudrun Kohl, Petra Pjevac and Joana Seneca Silva from the Joint Microbiome Facility of the Medical University of Vienna and the University of Vienna for assisting with amplicon and metagenomic sequencing, as well as repositing of sequencing data. We thank Sophia Derdak and Michael Schuster for initial data analysis. We thank Robert Vilvoi and Stephan Hemm for animal handling, Marcel Kertesz for mouse genotyping and Salwan Roumaia for NGS sample preparation. Treatment schemes and graphical abstract were created with BioRender.com.

This work was supported by the Austrian Science Fund, grant number ZK 57-B28 to CV, RG and FCP, grant number V 1025-B to RG, grant numbers DOC32-B28 to RV and MFJ and F8007 and P32678 to MFJ.

## Author contributions

RG, FCP, and CV: guarantor, supervision, conceptualization, methodology & experimentation, formal analysis, data curation, validation, visualization, resources. RG and CV: writing original draft and editing. MFJ: guarantor, supervision, conceptualization, resources and editing. RV, LC, MJ: conceptualization, methodology & experimentation. AH, KM, KDP, TvW, LS, MP, NI, IDV: methodology & experimentation. FCP, FD, AT, BH and NT: bioinformatics data curation, analysis and visualization. LB: resources. CG, DB, CB and MS: supervision. ML: ethics application for human study. CG: provided and cared for study patients.

## Online supplemental material

**Figure S1**

*FLNA editing impacts epithelial transcriptional profiles and inflammation in the healthy gut*.

**A)** Relative numbers of immune cells in healthy mouse colons (Student’s *t-*test). **B)** Gating strategy of colon tissue FACS analysis. Volcano plots (DEGs, padj ≤ 0.05) of FLNA^R^ versus FLNA^Q^ cells identified by DESeq2 analysis for **B)** EC progenitors, **C)** distal EC type II and **D)** goblet cells.

**Figure S2**

*FLNA editing protects mice from DSS-induced colitis and reduces early inflammatory cytokine production and intestinal neutrophil accumulation*.

**A)** Bodyweight of FLNA^WT^/IL-10^+/+^, FLNA^WT^/IL-10^−/−^, FLNA^Q^/IL-10^−/−^ and FLNA^R^/IL-10^−/−^ mice at the time of sacrifice (7-10 weeks of age, at onset of symptoms, 1-way ANOVA). **B)** Length measurements of colons at time of sacrifice (1-way ANOVA). **C)** Total colitis histology score. Higher score means more severe colitis. (Kruskal Wallis test, FLNA^WT^-IL-10^+/+^ and FLNA^WT^-IL10^−/−^, *n*=6; FLNA^Q^-IL10^−/−^, *n*=11; FLNA^R^-IL10^−/−^, *n*=12). **D)** Colon length measurements after five days DSS of FLNA^Q^ and FLNA^R^ mice (*n=*5 for each genotype, Student’s *t-*test). **E)** Cell clustering of large intestine mucosal cells after five days DSS. **F)** Absolute cell counts of immune cells from colons of mice treated with DSS for zero, two, five and ten days. **G-I)** Relative abundance (% of viable of CD45^+^ cells) of depicted immune cells of FLNA^R^ and FLNA^Q^ mouse colons at day two, five and ten of DSS treatment (1-way ANOVA). **J)** Gating strategy for flow cytometry of intestinal cells of FLNA^R^ and FLNA^Q^ mice. ***p<0.001; **p<0.01; *p<0.05. Error bars show SEM.

**Figure S3**

*A potentially protective microbiome does not explain DSS-resistance of FLNA^R^ mice*.

**A-B)** Pearson correlation analyses of ASV_1lf_am9 (belonging to the genus *Lachnospiraceae* NK4A136 group) and butyrate concentrations (A), or colitis score (B). **C)** Colony forming unit (CFU) determination after plating serial dilutions of fecal pellet slurries (*n=*4 for control and antibiotic (Abx) treated mice). **D)** Fecal microbial loads of control (*n*=4 FLNA^Q^, *n*=3 for FLNA^R^) and antibiotic treated (Abx) mice (open circles*, n*=3) as assessed by flow cytometry (Welch two sample *t*-test). **E)** Family-level relative abundance profiles of gut microbiomes from FLNA^R^ and FLNA^Q^ control or Abx treated mice. Families with relative abundances <2% are collapsed into the category “Other”. Each bar represents one mouse. **F)** Alpha diversity metric (Shannon Index) of samples from FLNA^Q^ and FLNA^R^ control and Abx treated mice, as assessed by 16S rRNA gene amplicon sequencing analyses (*n*=7 for Q Abx; *n*=8 for Q control, R control and R Abx; Welch two sample *t*-test). In D, E each point represents one mouse and boxes represent median, first and third quartile. Whiskers extend to the highest and lowest values that are within one and a half times the interquartile range. ****p<0.0001; ***p<0.001.

**Figure S4**

*Edited FLNA^R^ in myeloid cells protects from colitis*.

**A)** Sequencing chromatograms showing the validation of FLNA mutation status on RNA level (“mutant codon” = CAG, with either A or G in the centre): RT-PCR products sequenced around the *Flna* editing site (cttcAggga). FLNA^QiR^ mouse tissue (spleen) only expresses FLNA^Q^ (only “A” peak). A change from “A” to “G” (representing inosine (“I”) in the RNA) occurs only in Cre-recombinase positive hematopoietic cells (FLNA^QiR^ Vav-Cre, in spleen as a tissue enriched in hematopoietic cells, compared to colon and stomach tissues). **B)** Bodyweight loss curves of FLNA^QiR^ or FLNA^QiR^ Vav-Cre mice after DSS treatment until day ten (2-way ANOVA). **C-D)** Leukocyte counts eight weeks post transplantation of control (GFP+), FLNA^Q^ and FLNA^R^ BM into wildtype mice in bone marrow and blood (2-way ANOVA). **D-E)** Chimerism in cell populations eight weeks post transplantation of control GFP+ BM into GFP-wildtype mice in bone marrow and blood (2-way ANOVA). **G-H)** Bodyweight loss and total histopathological colitis score of wildtype mice transplanted with either FLNA^Q^ or FLNA^R^ BM and subsequently treated with DSS. Weights and scores shown at day ten (Student’s *t-*test and Mann Whitney U test). **I)** Bodyweight loss of FLNA^QiR^ or FLNA^QiR^ LysM-Cre mice after DSS treatment at day ten (Student’s *t-*test). **J)** Scheme for CD8+ T cell depletion and DSS challenge. **K)** Colon length measurement of CD8+ T cell depleted FLNA^Q^ and FLNA^R^ mice at day ten (Student’s *t-*test); **L)** Bodyweight loss curves until day ten (2-way ANOVA); and **M)** Total histopathological colitis score at day ten of DSS challenge (Mann Whitney U test). Arrows in L indicate timepoints of antibody injections. *p<0.05. Error bars show SEM.

**Figure S5**

*Edited FLNA^R^ in myeloid cells protects from colitis*.

**A-F)** Correlations of delta C_T_ values from RT-qPCR analysis of mRNA expression versus FLNA editing levels determined by amplicon sequencing analysis in distal colon on days: zero, two, five, and ten (Simple regression analysis). Higher dC_T_ = lower expression. **G)** Dot plots representing *Adar* and *Adarb1* expression in log counts per million (log(CMP+1) in untreated FLNA^Q^ and FLNA^R^ mouse colons determined by scRNA-Seq in different cell populations. **H)** Expression levels of *Adar* and *Adarb1* in WT colon cells analyzed by scRNA-Seq and shown as heat-gradient in the clustered cells (log(CMP+1). **I)** Representative images of colon histology sections stained for NETosis (by CitH4 AB - red) and Neutrophils (by Ly-6G AB - green) of at day ten of DSS in: FLNA^Q^ (left) and FLNA^R^ (right); DAPI in blue. Scale bar = 100µm. **J-K)** Selected interactions quantified by CellChat analysis of scRNAseq data of FLNA^Q^ and FLNA^R^ mouse colons post five days of DSS challenge. Communication probabilities (interaction strength) of different cell types to neutrophils (J) and monocytes/macrophages (K).

