## Supplementary figures and images for "Edited Filamin A in myeloid cells reduces intestinal inflammation and protects from colitis"

### Supplementary Figure 1

Supplementary Figure1

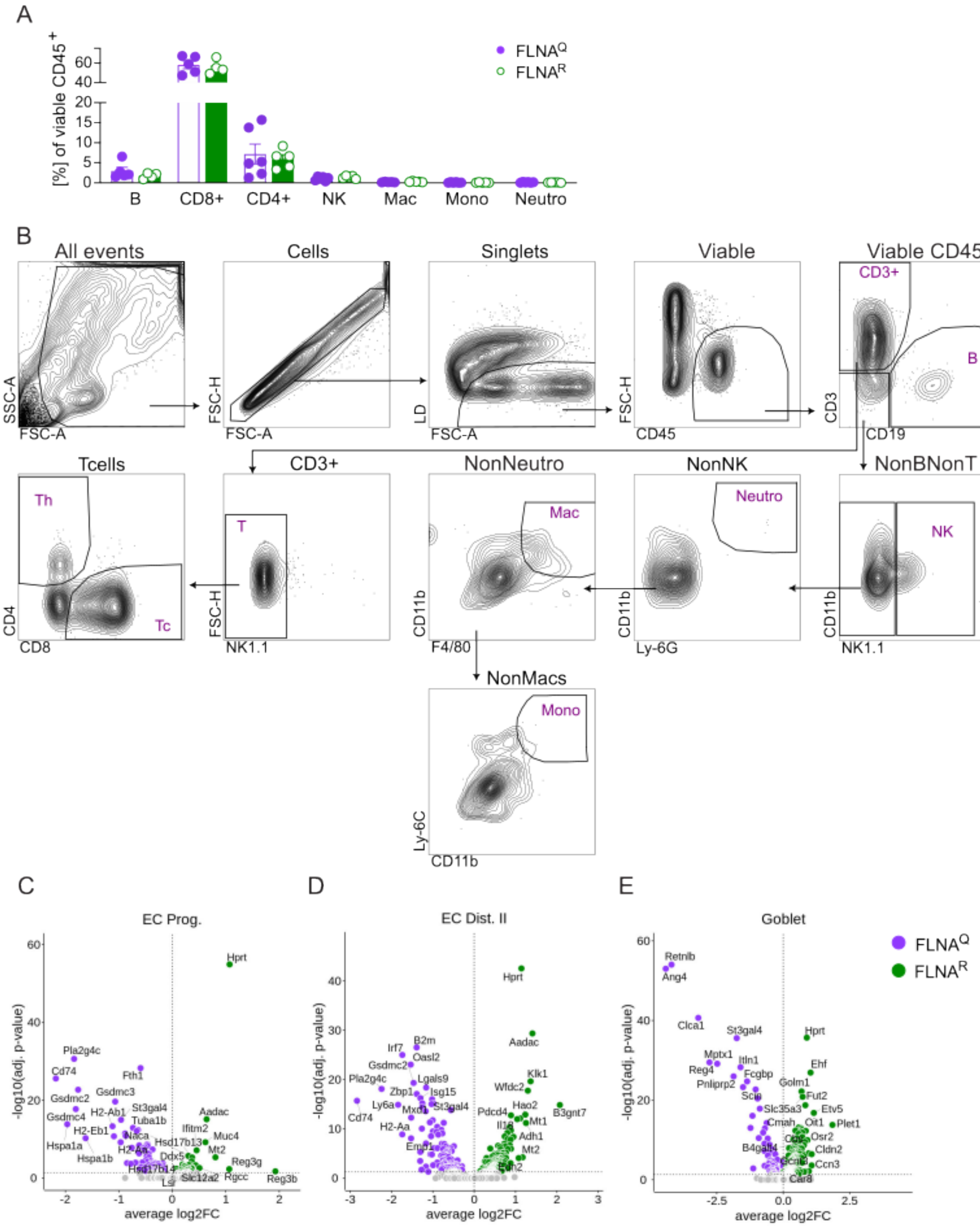

### Supplementary Figure 2

#### Supplementary Figure 2

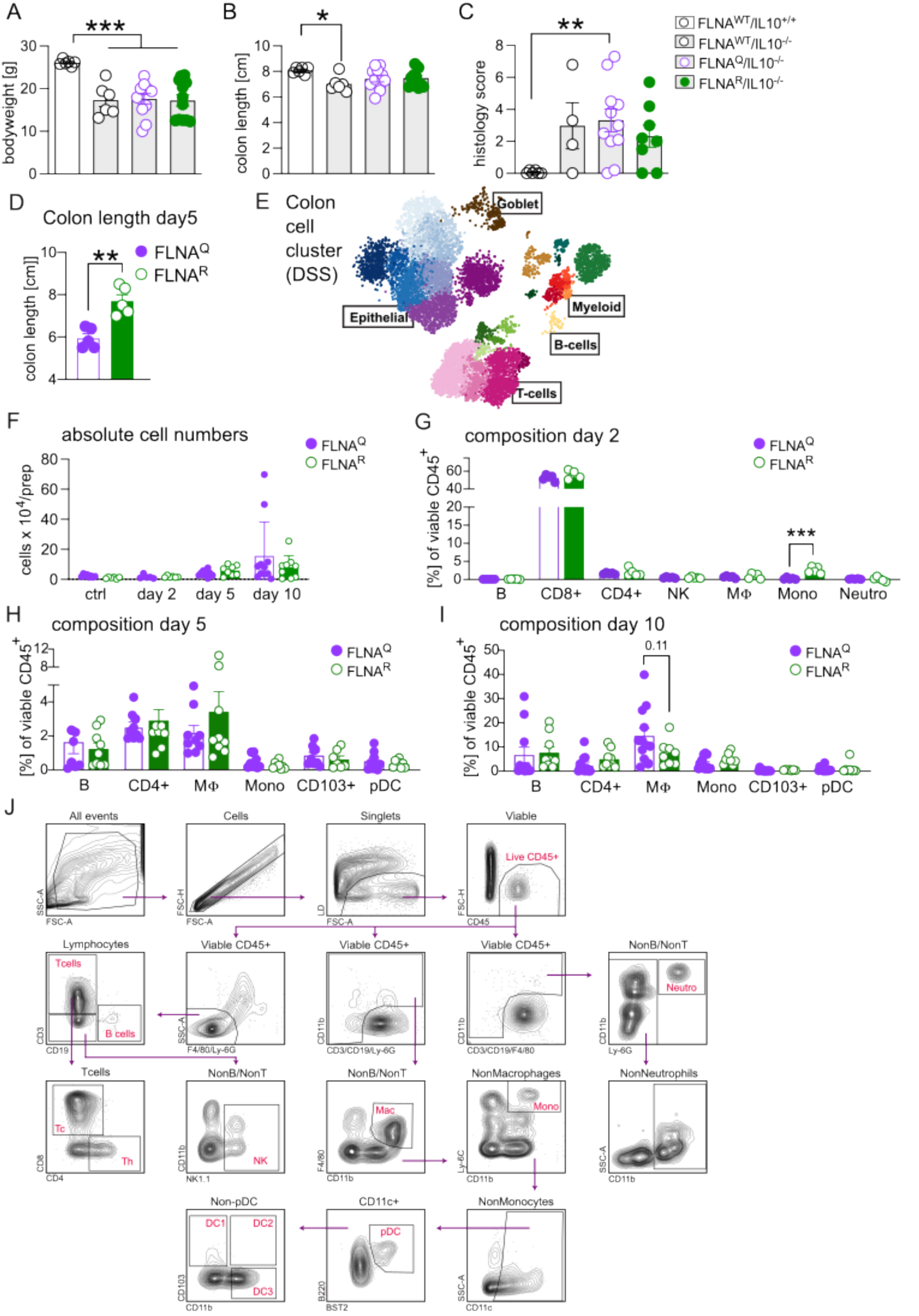

### Supplementary Figure 3

Supplementary Figure 3

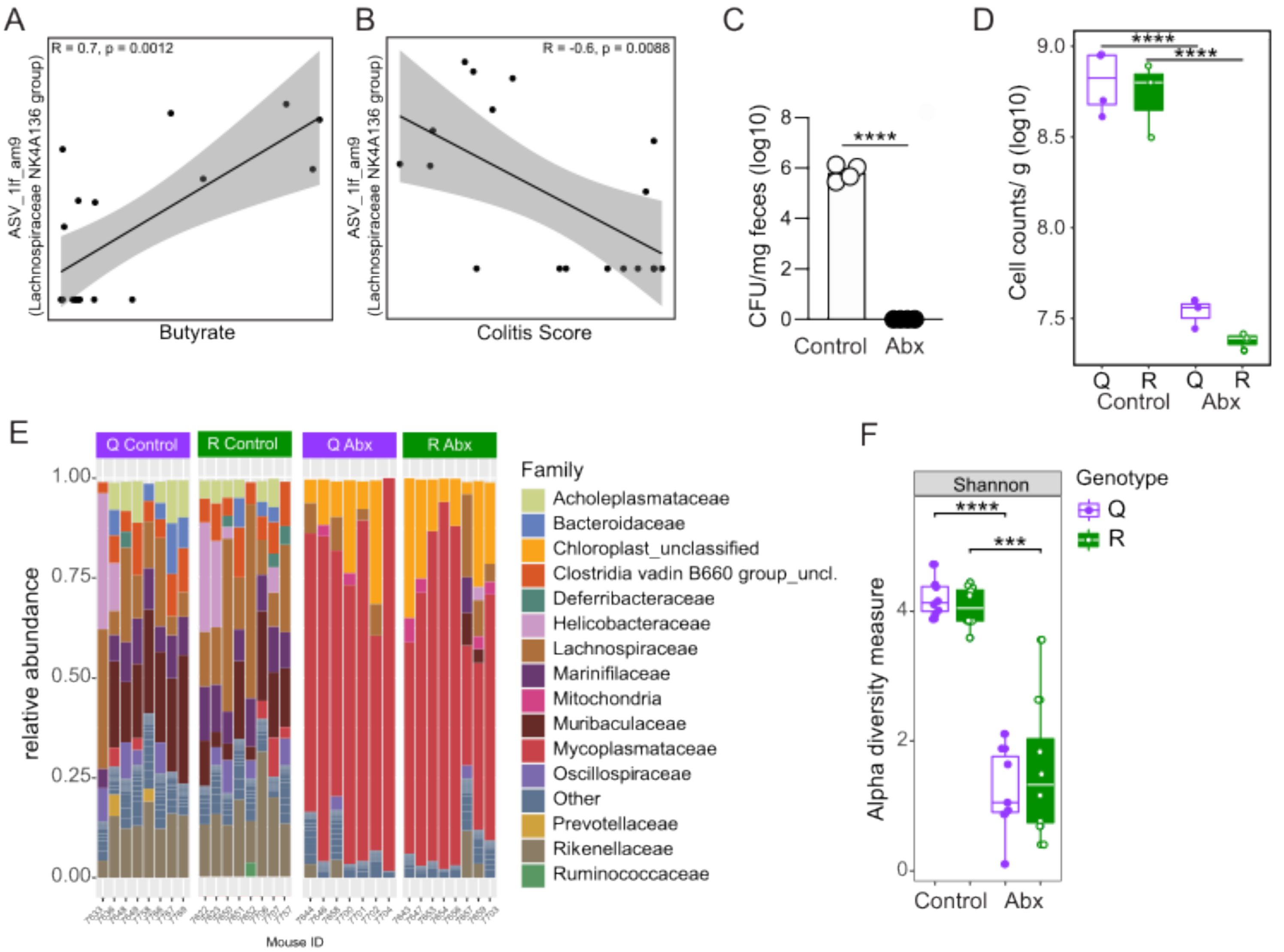

### Supplementary Figure 4

Supplementary Figure 4

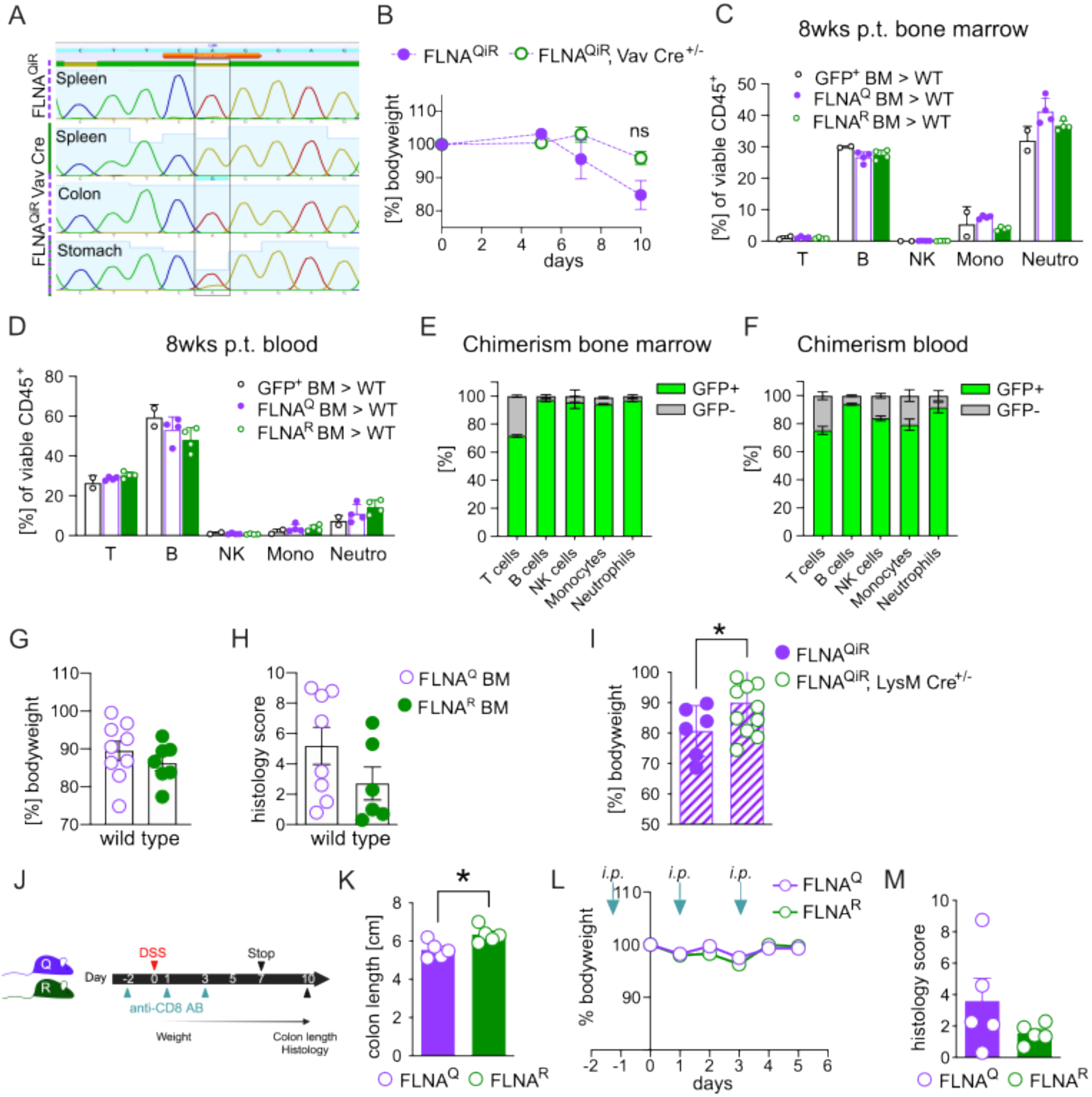

### Supplementary Figure 5

Supplementary Figure 5

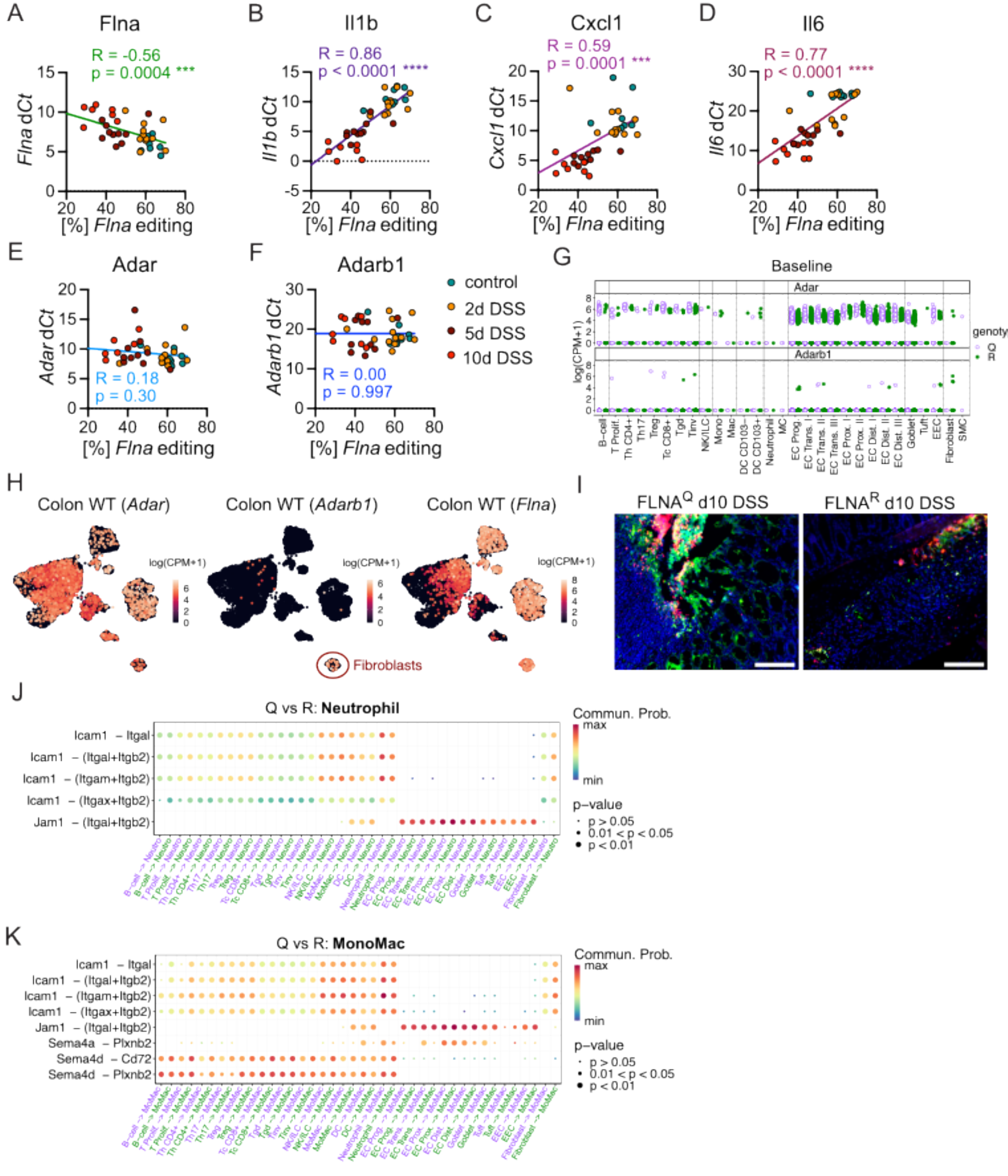
